# Cerebello-Basal Ganglia Functional Network Integration in Psychosis

**DOI:** 10.1101/2022.11.12.516285

**Authors:** T. Bryan Jackson, Katherine S. F. Damme, Vijay A. Mittal, Jessica A. Bernard

## Abstract

Psychotic disorders are conceptualized as brain-network diseases and both the cerebellum (CB) and basal ganglia (BG) are implicated in widely used conceptual models. Previous research has focused on these structures and their respective circuits as distinct, however, both are functionally and anatomically connected to each other and to cortical networks via domain-specific, topographically organized thalamo-cortical loops. Currently, it is unclear how CB-BG network dysfunction may play a mechanistic role in the course of psychosis; however, network global efficiency (GE), a measure of functional integration, is a novel approach that aims to represent cognitive and motor CB-BG network (CCBN, MCBN, respectively) connectivity in cross-sectional groups of healthy control (HC), clinical high-risk (CHR), early course psychosis (ECP), and chronic psychosis (CP) participants. We compared network GE between groups and inspected individual differences in CCBN- and MCBN-GE as it relates to group membership and to psychosis symptoms. We also associated CB-BG network GE with cortical network GE. Results indicated that CCBN-GE was associated with cognitive dysfunction and lower in CHR individuals, compared to HC and CP; while MCBN was associated with negative psychosis symptoms. Last, we detailed CB-BG associations with sensory, motor, default mode, and salience networks across groups, with group effects demonstrating complex differences within the ECP group. Findings indicating that CB-BG network dysfunction may play an important role in early pathogenesis and authors argue for CB-BG dysfunction to be analyzed from a network perspective. Future work is needed however to incorporate this approach into our understanding of psychosis.

## Introduction

Decades of evidence including increased striatal dopamine^1–4^ and volumetric differences within the caudate and striatum^5,6^ have implicated basal ganglia (BG) dysfunction in psychotic disorders. A study investigating medicated patients with schizophrenia found larger bilateral striatal volume^7^, and a more recent work has suggested links between basal ganglia function and negative symptomatology^8^. Decreased functional activation within BG structures has also been noted in patients with schizophrenia^9,10^. Additionally, conversion to a psychotic disorder in clinical high-risk (CHR) samples was predicted by motor deficits related to BG dysfunction^11,12^. In short, the BG are of particular importance when discussing psychotic disorders and symptomology.

In parallel, the cerebellum (CB) has also been linked to psychotic disorders and their symptoms. For example, differences in the CB were linked to psychosis-related disorders due to its role in working memory, language processing, and general cognition, leading researchers to propose the term “cognitive dysmetria,” suggesting the cognitive deficits are similar in nature to the deficits seen in CB-related motor dysmetria^5,13–17^. Additionally, anhedonia was related to differences in CB network properties^18,19^. Meta-analytic evidence has also supported a role for the CB in psychosis. This includes work that pointed to differences in CB structure and functional connectivity during the first-episode of psychosis, including smaller CB Crus I and Crus II volume^20^, and that which found differences in regional activation patterns during emotional, cognitive, and motor tasks, with Lobule VI and Crus II displaying consistently lower activation in patients with psychsis relative to controls^21^. Clearly subcortical dysfunction associated with psychotic disorders extends beyond the BG to include the CB as well.

Research has largely focused on BG and CB regions separately despite the preponderance of evidence suggesting that both structures are anatomically and functionally altered in psychosis. Increasingly, the BG-centric view has shifted to encompass additional regions and psychotic disorders are now largely seen as disorders related to network dysfunction^22^. Indeed, the CB, BG, and their thalamo-cortical networks have each been implicated in many psychiatric and neurological disorders, including schizophrenia and other psychosis-related disorders^23–27^. For example, reduced functional connectivity between the CB and cortical networks in patient groups has been noted^28^ and patient groups have decreased CB and BG global efficiency (GE)^22,29^.

There are also network differences related to symptomology. Dynamic functional connectivity between the CB and the fronto-parietal network coupled with grey matter volume has been associated with positive symptoms^28,30^ and those with schizophrenia do not show a relationship between cerebello-cortical connectivity and processing speed (as opposed to healthy controls [HC] who do)^31^. Additionally, higher functional connectivity (FC) between the Lobule V and the motor cortex was associated with worse positive symptoms 12 months later while FC between Crus I/Crus II and the cingulate cortex marginally predicted positive symptom change in CHR individuals^32^. Of note, one study found schizophrenia patients had both increased FC between bilateral caudate nucleus and decreased FC between the BG and Crus I/Crus II of the CB (regions associated with cognition and working memory)^33^, supporting the use of CB-BG network integration metrics in determining biomarkers for psychosis symptomology.

Connectivity studies have provided a framework for understanding interactions between CB-BG networks. The regions are highly functionally^34,35^ and anatomically^36,37^ connected via disynaptic subcortical circuits and communicate with the cortex (and indirectly with each other) via thalamo-cortical loops^38,39^. Alterations in these cortical pathways are associated with psychosis risk ^40^ and age of onset^41^. Additionally, we reported positive associations between the integration of CB-BG networks and that of cortical networks in HC^35^ and found that motor CB-BG network (MCBN) GE negatively predicted self-reported depression and hyperactivity symptoms in an unselected community sample^42^.

To this point, no known study has investigated differences in CB-BG network integration as it relates to psychotic symptoms or cortical network integration in psychosis patients. Yet, given that CB-BG networks support many cognitive, motor, and limbic processes^43^, the noted psychosis-related associations, and the high interregional connectivity in healthy young adults^35^, differences in CB-BG network connectivity may help explain symptom differences in CHR, early course psychosis (ECP), and chronic psychosis (CP) patients. Therefore, research into CB-BG network differences during the development and progression of psychosis may be important for predicting disease onset and developing new targets for intervention, thereby improving health and quality of life for individuals at risk for psychosis. In this study, we used a novel approach to investigate differences in functional network integration (network global efficiency, GE). We measured GE within cognitive and motor CB-BG networks (CCBN and MCBN, respectively) and determined whether these measures were associated with psychosis symptoms and diagnoses within a sample of healthy controls, individuals diagnosed with psychosis spectrum illnesses, or noted as being CHR.

Here, we combined subject data from three sources to create a sample that captures the disease progression from CHR, to ECP, and finally to CP patients. We measured the GE of the CCBN and MCBN and assessed the differences between groups and relationships with psychosis symptomology across groups. Given that connectivity within and between the CB and BG have been investigated in the past, we predicted reduced GE in patients^30,33^ and an association with cognitive dysfunction^30,44^. We also predicted an association between CB-BG GE and both positive^28,32^ and negative symptoms^8^. Lastly, we assessed the relationships between CB-BG and cortical network GE across groups using group as a predictor. We expected relationships between GE of CBGN and default mode (DMN) and auditory network (AN), and between MCBN and cingulo-operculum (CON), motor (MN), visual (VN) and AN^35,42^.

## Method

### Participants

Our final sample consisted of 448 participants from three different datasets: 1) A private dataset (n = 172) collected by the Adolescent Development and Prevention (ADAPT) research lab at Northwestern University that included CHR and healthy control participants^9,21,45,46^; 2) a publicly-available dataset collected by the Center for Biomedical Research Excellence (COBRE) and made available via schizconnect.org^47,48^ that included a broad range of psychosis patients and healthy controls of all ages (n = 153); and 3) a publicly available dataset collected by the Human Connectome Project for Early Psychosis (HCP-EP) that included early course psychosis patients (defined as within 5 years of a psychosis diagnosis) and healthy controls (n = 123). Participants that had both resting-state and anatomical images and symptom scores were included. 9 participants (1 CHR, 6 CP, 2 HC) were removed due to having no within-network connectivity for any network, indicating issues with data quality. Table 1 details relevant demographic information.

**Table 1.** Descriptive statistics. Mean (standard deviation).

|  | HC (n = 204) | CHR (n = 91) | ECP (n = 104) | CP (n = 49) |
| --- | --- | --- | --- | --- |
| Age | 27.7(12.4) | 19 (2.12) | 22.71(2.49) | 43.6 (10.45) |
| Alcohol (lifetime) | .79 (.79) | .40 (.49) | 1.23 (.62) | 1.37 (.67) |
| Marijuana (lifetime) | .80 (.86) | .1 (.34) | 1.52 (.86) | 1.37 (.70) |
| CPZ Equivalency | 0 | 0 | 411 (648) | 1160 (1400) |
| PANSS-P | 4.16 (6.42) | - | 15.1 (5.12) | 15.1 (4.07) |
| PANSS-N | 3.68 (5.94) | - | 14.1 (5.92) | 15.0 (5.30) |
| PANSS-C | 9.07 (2.68) | - | 11.43 (3.28) | - |
| SIPS-P | .60 (1.27) | 11.98 (4.46) | - | - |
| SIPS-N | .44 (.76) | 8.44 (6.35) | - | - |

PANSS – Positive and Negative Symptom Scales; SIPS – Structured Interview for Prodromal Symptoms; P – positive score, N – negative score, C- cognitive score. Lifetime marijuana and alcohol usage were assessed using the Structured Clinical Interview for Disorders IV and V (see “Group Construction” for additional details about which version was used).

### Symptom Measures

CHR and HC participants within the ADAPT dataset were interviewed and scored using the Structured Interview for Psychosis-risk Syndromes (SIPS)^49^. ECP, CP, and HC participants from the COBRE and HCP-EP participants were assessed using the Positive And Negative Syndrome Scale (PANSS)^50^. The HCP-EP dataset also contained a cognitive subscale of the PANSS and was collected using the same methods within that dataset only. Table 1 details the mean and standard deviation of the symptoms scores.

### Anti-psychotic Medication

Some ECP and CP patients were medicated using one or more anti-psychotic medication. All anti-psychotic medications were converted to a common metric, chlorpromazine (CPZ) equivalency (Table 1), using published equivalencies for oral conventional and atypical antipsychotics^51^. To account for differences in FC due to anti-psychotic medication, CPZ equivalency was entered into the regressions as a covariate.

### Group Construction

Data were concatenated across datasets and sorted into 4 groups based on diagnoses and time since diagnosis. ADAPT and COBRE participants underwent the Structured Clinical Interview for Axis-I DSM IV Disorders^52^; HCP-EP participants underwent the Structured Clinical Interview for Axis-I DSM V Disorders^53^ (SCID). HC had no history of a psychotic disorder nor a first-degree family member with a psychosis or bipolar disorder diagnosis, no history of major depression, and were not taking any psychiatric medications. Scores for lifetime marijuana and alcohol use were measured during the structured interviews and included in analyses as a covariate. HC participants were combined from all datasets into one pool (n = 204). Inclusion in the CHR group (n=91) was determined by having no Axis-I disorder, moderate levels of positive symptoms (a SIPS score of 3–5 in one or more of the 5 positive symptom categories), and/or a decline in global functioning in association with the presence of schizotypal personality disorder, and/or a family history of schizophrenia, consistent with prior investigations using this dataset^21,45,46^. Inclusion as an ECP patient for the HCP-EP dataset (n = 80) was determined by a psychosis diagnosis (schizophrenia, schizophreniform, schizoaffective, delusional disorder, brief psychotic disorder, or psychosis not otherwise specified) determined by completing a SCID, and by time since diagnosis (5 or fewer years with the psychosis diagnosis). Inclusion as an ECP or CP patient for the COBRE dataset was determined by a diagnosis of schizophrenia or schizoaffective disorders, determined by completing a SCID. We wanted to capture differences between ECP and CP patients but did not have access to time since diagnosis information for users in the COBRE dataset. Given the average age of diagnosis is early adulthood,^54^ we assumed there would be some in the COBRE dataset that were similarly in the early stages of psychosis treatment. To account for these individuals, we age-matched COBRE patients (n = 24) to patients from the HCP-EP dataset to create the ECP group (n = 104). Remaining patients in the COBRE dataset were assigned to the CP group (n = 49).

### Data Acquisition and Preprocessing

Raw, unpreprocessed functional and anatomical data were downloaded with the consent of the custodian of each dataset. Results included in this manuscript come from preprocessing performed using fMRIPrep 21.0.0^55,56^, which is based on Nipype 1.6.1^55,57^. A full description of the preprocessing steps taken directly from fMRIPrep’s boilerplate output is included in Supplementary Material 1 for replication purposes. Of note, the ADAPT dataset did not contain fieldmaps or oppositely encoded images. While suboptimal, we prioritized standardization of the combined dataset to reduce between-dataset variance introduced due to preprocessing and thus did not perform any unwarping steps on the functional data for any dataset. Following preprocessing, structural images underwent tissue type segmentation and functional images underwent smoothing (5 mm FWHM), artifact detection (global signal z-value threshold: 5, subject motion threshold: 0.9 mm), and denoising using the CONN toolbox (v. 21a)^58^. Please refer to the Supplementary Materials 1 for a full description.

### Regions of Interest

Using fslmaths, 3.5 mm spherical, binarized regions of interest (ROIs) representing the CCBN and MCBN and 7 rsfMRI cortical networks, described below, were created with coordinates from previous works (FSL v.6.0.3: FMRIB’s Software Library)^59^. Supplementary Table 1 details the 122 resulting ROIs. CB-BG network ROIs were created using CB and BG coordinates previously used in recent work in our lab^34,35^. Lobule V and Lobule VI of the CB and dorsal caudal and dorsal rostral putamen of the BG were used to define the MCBN; while Crus I and Crus II of the CB and the inferior and superior ventral striatum, dorsal caudate, and ventral rostral putamen of the BG were used to define the CCBN^34,35^.

We similarly constructed cortical networks spanning cognitive (fronto-parietal [FPN], CON, and DMN), motor (MN), emotional (EN), and sensory (VN and AN) domains. Cortical networks were defined using ROIs from previous reports^60,61,62,63,64^. Coordinates and region names are included in Supplementary Table 1 and fully detailed in our previous work^35^.

### First-level fMRI Analysis, Global Efficiency

All participant-level computations were performed in the CONN toolbox (v. 21a)^58^. For each participant, ROI timeseries were extracted and cross-correlated using bivariate analyses, resulting in a 122×122 correlation matrix for each participant. To compute graph theory components, edges were defined as thresholded correlation coefficients and nodes were defined as the ROIs in each network as described above^34,60,62,65^. We previously investigated differences in network measures when different thresholds were^35^ used and found no substantial differences; as such, we used *β* > .1 to define edges, a correlation coefficient threshold commonly used in the literature^60^ and consistent with our prior work^35^. To define interconnectedness, network-level GE, a graph theory metric of how efficiently information in a network travels, was computed separately for each of the nine networks^58,66–68^.

### Analysis Design

All analyses were conducted in RStudio (version 2022.02.2; rstudio.com) using base R and the stats package (v 4.2.0)^69^. An RMarkdown file including the code used and their outputs are provided in Supplementary Material 2. ANCOVAs were preformed using the “aov” command and regression models using the “lm” command. First, we used ANCOVAs to describe group differences in CCBN GE, MCBN GE and cortical GE. The models controlled for lifetime alcohol and marijuana usage, age (assessed using SCID-IV and SCID-V), dataset membership, and CPZ equivalency. Second, we used ANCOVAs to describe group differences in psychotic symptomology, controlling for alcohol and marijuana abuse, age, and CPZ equivalency. All ANCOVAs were followed up using t-tests comparing each group pair. Analyses and FDR correction were performed using the glht function from the multcomp package (v.4-19)^70^.

Next, we used standard regression to probe whether CCBN and MCBN GE predicted positive (PANSS-P, SIPS-P), negative (PANSS-N, SIPS-N) and cognitive (PANSS-Cog) symptomology, controlling for alcohol, marijuana, age, and CPZ equivalency. A separate regression model was computed for each predictive measure which was necessary given that some measures were limited to only certain subset of the data.

Finally, we conceptually replicated our previous work that related CB-BG and cortical network GE. We modified our analyses by entering CCBN GE, MCBN GE, group (as a factor), and their interactions as predictors for cortical GE, controlling for alcohol, marijuana, age, dataset membership, and CPZ equivalency. We followed up significant group effects and group interactions by performing the same ANCOVA within the affected group.

### Results Group Differences in Symptomology and CB-BG GE

Significant group differences were found in all psychosis measures in the expected directions (*ps* < .001; Figure 1A), indicating CHR individuals have higher scores than HC and a potential increase in psychotic symptomology from ECP to CP.

**Figure 1:**
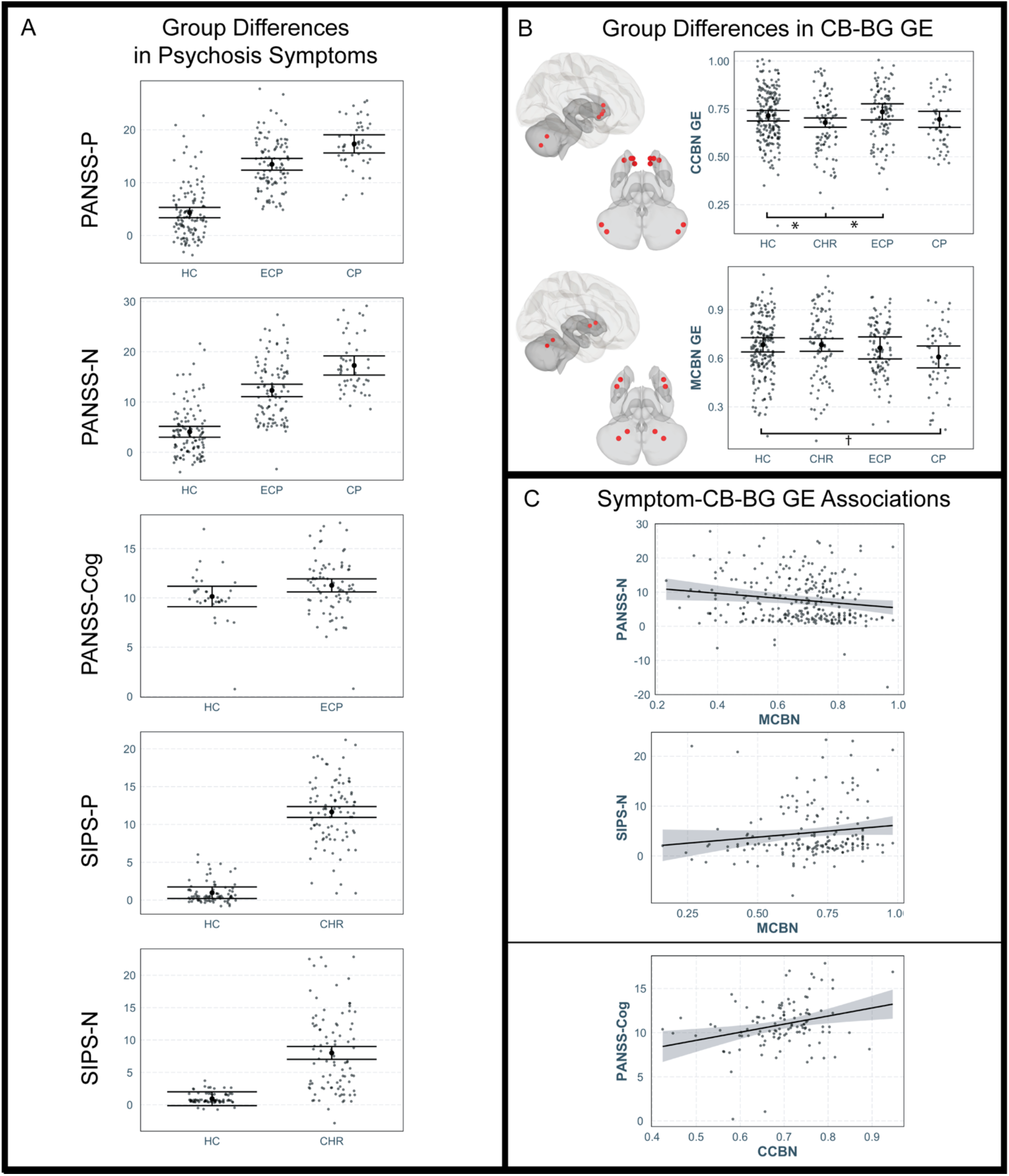
A: group differences in symptomology. B: cerebellum (CB) and basal ganglia (BG) ROIs overlayed on MNI152 template and group differences in CB-BG global efficiency (GE). C: Scatterplots depicting associations between CB-BG GE and symptomology.

Group differences in CCBN GE (Figure 1B) indicated a significant main effect of group (*F*(3, 428) = 2.91, *p* = .03) and post-hoc tests indicated CCBN GE was significantly higher in ECP (*t*(194) = 2.67, *p* = .03) and HC (*t*(294) = 2.54, *p* = .03) compared to CHR. Group differences in MCBN GE (Figure 1B) indicated a significant main effect of group (*F*(3, 428) = 3.97, *p* = .01) and post-hoc tests indicated MCBN GE was marginally lower in CP compared to HC (*t*(252) = 2.47, *p* = .08).

### CB-BG Networks and Symptomology

All regression models predicting psychosis symptomology from CCBN and MCBN GE were significant (*ps* < .001; Figure 1C, Table 2). CCBN GE significantly predicted Cognitive PANSS scores within the HCP-EP ECP participants (*n* = 80; *p* < .01). MCBN GE significantly predicted both PANSS-N across HC, ECP, and CP groups collected from the HCP-EP and COBRE datasets (*p* = .02) and SIPS-N across the CHR and HC groups collected from the ADAPT dataset (*p* < .001).

**Table 2.** Statistics from the regressions predicting symptom scores. Each column presented represents a separate model predicting the measure. Beta values and (standard error) are displayed for each independent variable and control measure.

|  | PANSS-P | PANSS-N | PANSS-Cog | SIPS-P | SIPS-N |
| --- | --- | --- | --- | --- | --- |
| (Intercept) | 11.31 * | 16.56 *** | 5.03 | -1.58 | -0.28 |
|  | (4.43) | (4.78) | (2.85) | (5.57) | (5.30) |
| CCBN | -3.15 | -4.51 | 9.17 ** | -6.32 | -5.27 |
|  | (4.91) | (5.30) | (3.13) | (4.55) | (4.33) |
| MCBN | -3.90 | -7.12 * | -2.31 | 3.13 | 4.84 |
|  | (2.84) | (3.07) | (1.97) | (3.07) | (2.92) |
| Alcohol | 0.12 | -0.65 | -0.09 | 4.23 *** | 3.71 *** |
|  | (0.65) | (0.70) | (0.40) | (1.04) | (0.99) |
| Marijuana | 0.77 | 0.68 | 0.02 | 7.47 *** | 7.31 *** |
|  | (0.56) | (0.61) | (0.30) | (1.77) | (1.68) |
| Age | -0.05 | -0.07 | 0.01 | 0.47 * | 0.21 |
|  | (0.04) | (0.04) | (0.07) | (0.19) | (0.19) |
| Dataset HCP | 6.59 ***<br>(1.07) | 4.46 ***<br>(1.16) |  |  |  |
| CPZ_equivalency | 0.00 ***<br>(0.00) | 0.00 ***<br>(0.00) | 0.00 ***<br>(0.00) |  |  |
| N | 267 | 267 | 114 | 171 | 171 |
| R2 | 0.37 | 0.29 | 0.24 | 0.25 | 0.21 |

PANSS – Positive and Negative Symptom Scales; SIPS – Structured Interview for Prodromal Symptoms; P – positive score, N – negative score, C- cognitive score. CCBN – cognitive cerebellar-basal ganglia network, MCBN – motor cerebellar-basal ganglia network. \*\*\* $p < 0.001$ ; \*\* $p < 0.01$ ; \* $p < 0.05$ .

### Group Differences in Cortical GE

Group differences in cortical network GE (Figure 2A) were found in CON (*F*(3, 428) = 3.81, *p* = .01), DMN (*F*(3, 428) = 4.9, *p* < .01), EN (*F*(3, 428) = 5.45, *p* = .001), AN (*F*(3, 428) = 4.43, *p* < .01), and VN (*F*(3, 428) = 3.34, *p* = .02) GE, but no differences were found between groups in pairwise post hoc tests (*p’s* > .12).

**Figure 2:**
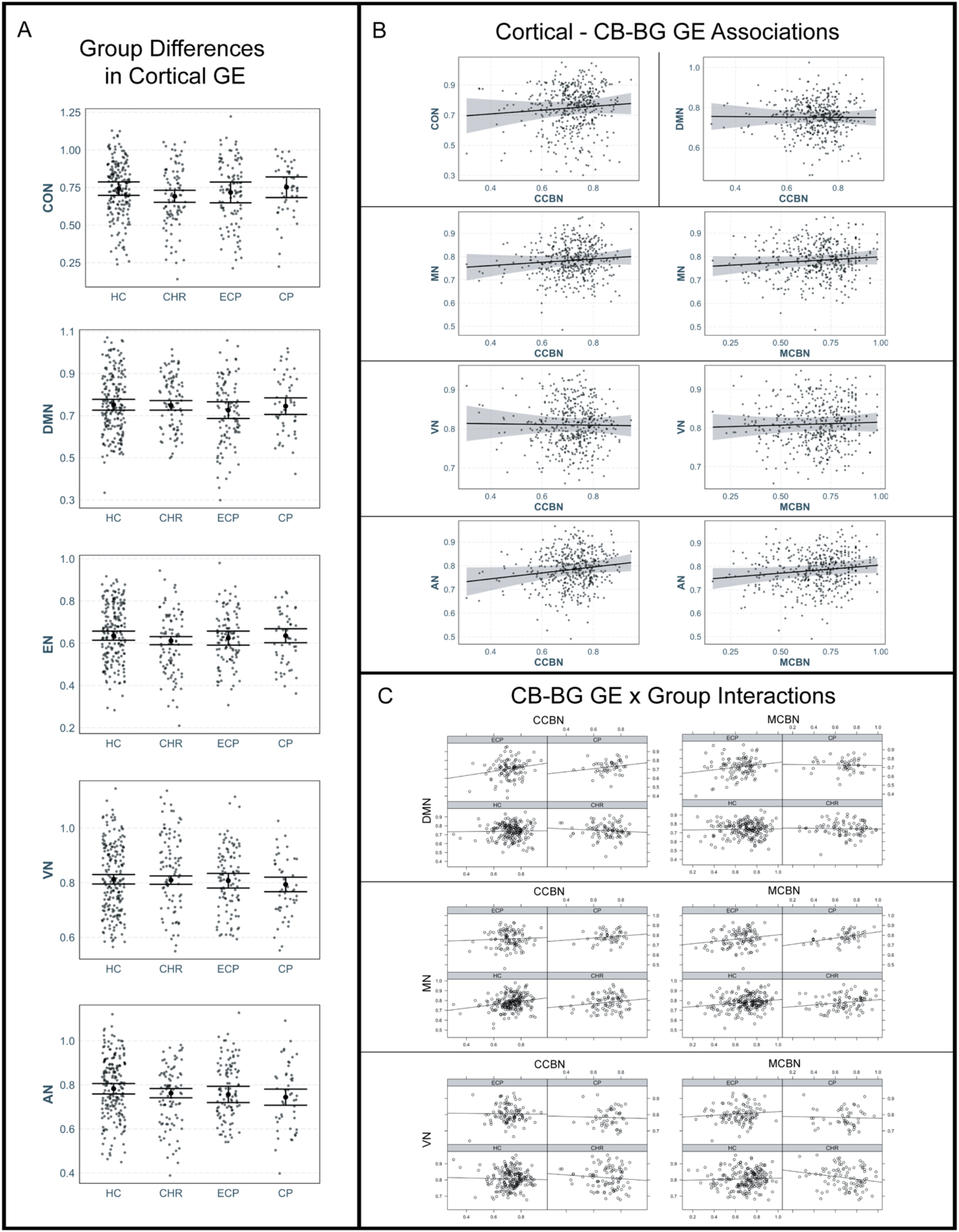
A: group differences in cortical global efficiency (GE). B: Scatterplots depicting associations between cortical and cerebellar-basal ganglia (CB-BG) GE across groups. C: Scatterplots depicting group x CB-BG GE interactions.

### CB-BG and Cortical Network Relationships Across Groups

Figures 2B and 2C and Tables 2 and 3 detail the following CB-BG – cortical network relationships. Three-way interactions between CCBN GE, MCBN GE, and ECP group membership significantly predicted MN GE (*p* = .02), and VN GE (*p* < .01) and marginally predicted DMN GE (*p* = .07), indicating CCBN and MCBN GE interacted within the ECP group. Two-way interactions between CCBN GE and ECP group membership significantly predicted DMN GE (*p* = .04), MN GE (*p* = .04), and VN GE (*p* = .002) and two-way interactions between MCBN GE and ECP group membership significantly predicted MN GE (*p* = .03) and VN GE (*p* = .03) and marginally predicted DMN GE (*p* = .06), indicating differences in CB-BG GE within the ECP group. An interaction between CCBN and MCBN GE marginally predicted CON GE (*p* = .06). CCBN GE significantly predicted CON GE (*p* = .03), EN GE (*p* < .001), and AN (*p* < .001). MCBN GE significantly predicted AN GE (*p* = .04).

**Table 3.**
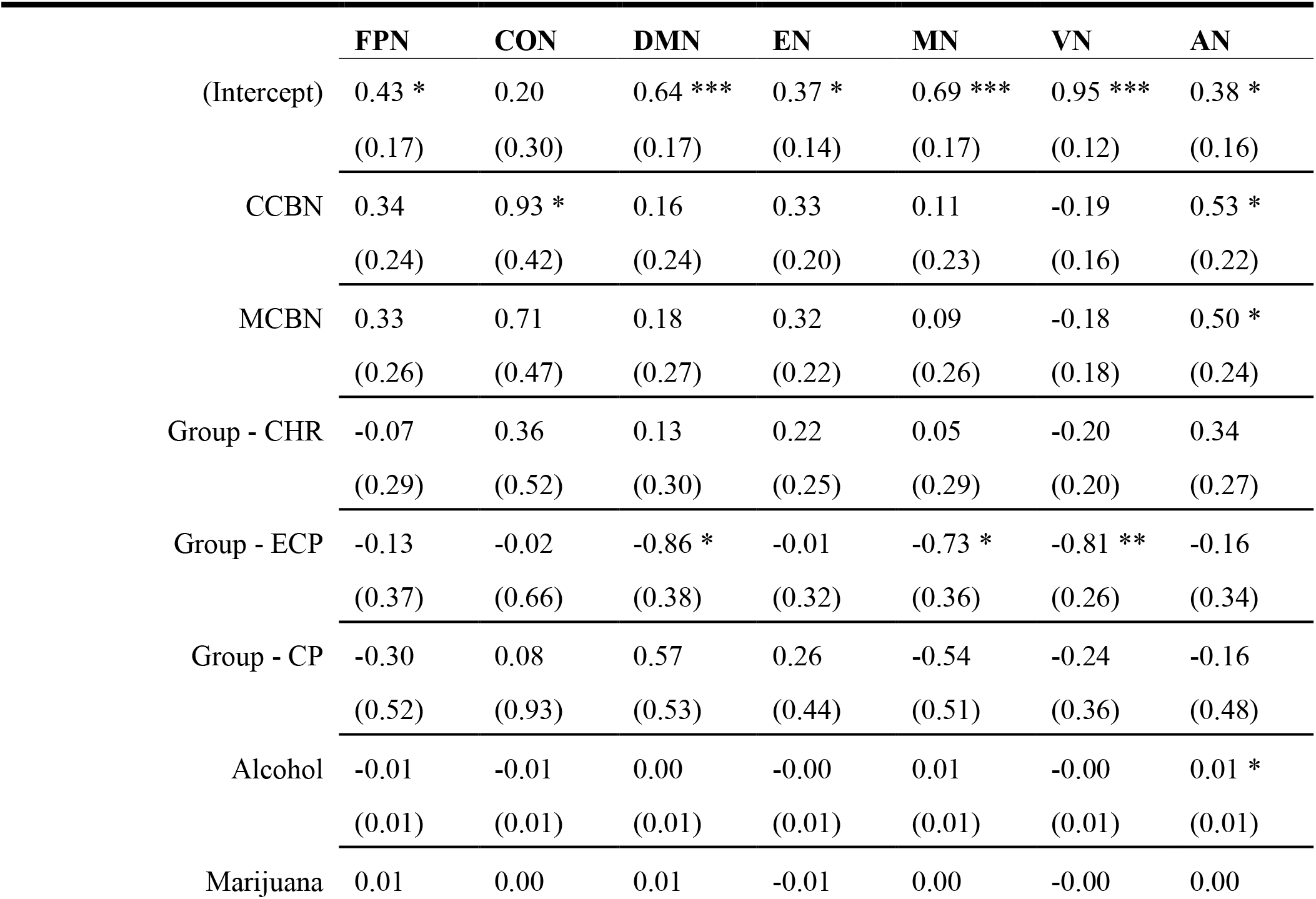
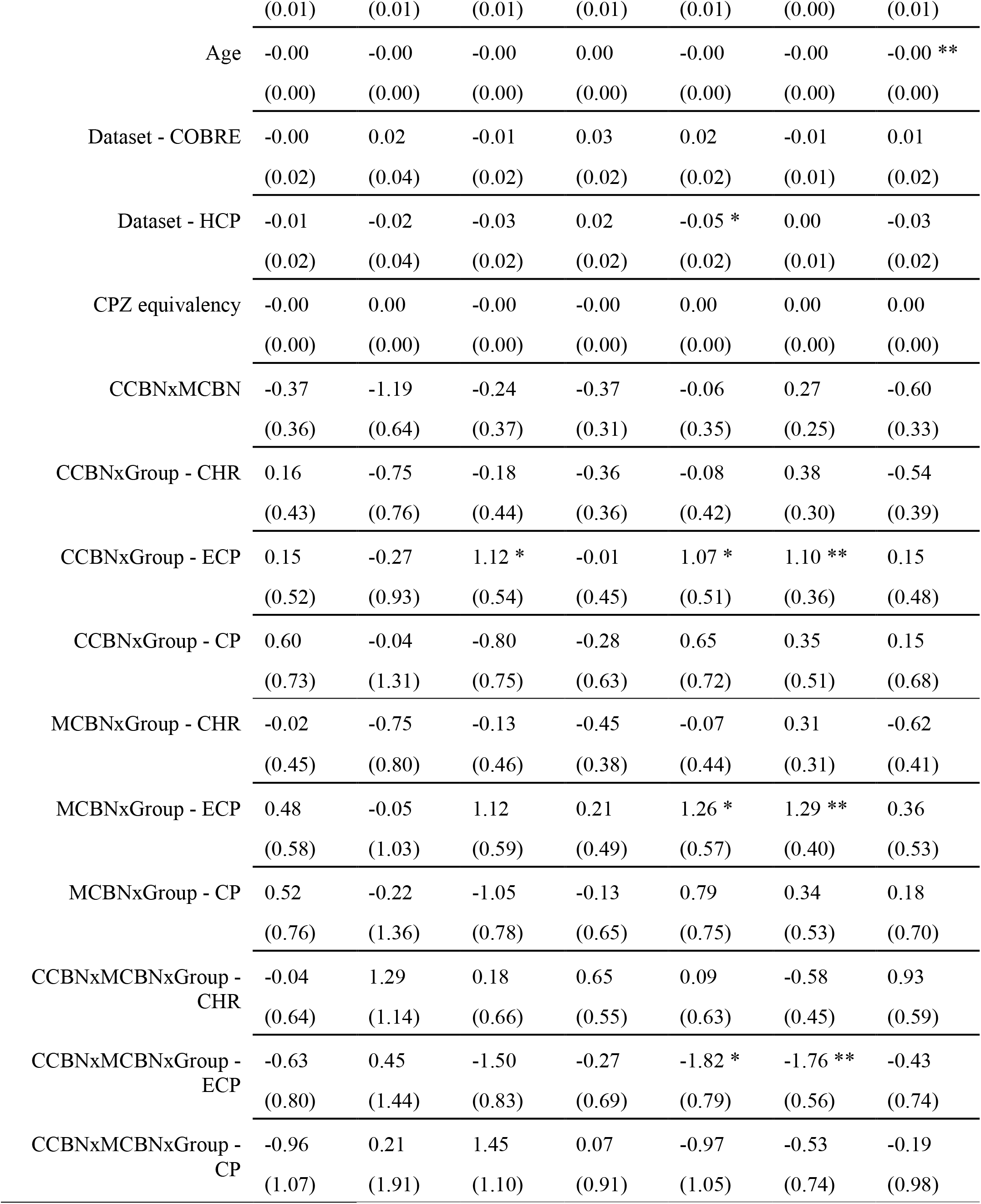
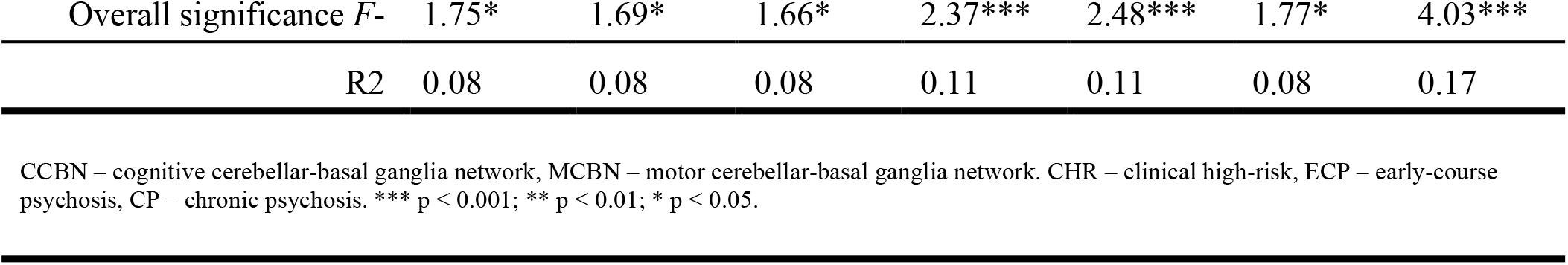
Statistics from the regressions predicting cortical global efficiency (GE) from cerebellar-basal ganglia GE. Beta values and (standard error) are displayed for each independent variable and control measure.

Post-hoc tests within the ECP group were performed to further investigate the interactions (Table 3). Models predicting MN (*p* = .02) and VN (*p* = .02) within the ECP group were significant. Interactions between CCBN and MCBN GE were significant for both MN (*p* = .01) and VN GE (*p* < .001) models, indicating non-additive effects of CB-BG GE. Both CCBN GE and MCBN GE were significant predictors in both models (*ps* < .01). The model predicting DMN GE from CB-BG GE within the ECP group was not significant, nor were any of the individual predictors within the model.

## Discussion

Here, we investigated CB-BG networks in those at risk for or diagnosed with a psychotic disorder and revealed cross-sectional differences in CB-BG network efficiency related to psychosis symptomology. Group differences in psychosis symptoms were all significant and in the expected direction. The CP group displayed higher positive and negative scores compared with the ECP group, consistent with findings suggesting increased scores with each incidence of symptom relapse^71^. Consistent with our hypotheses, we found that high CCBN GE predicted high cognitive dysfunction scores and low MCBN GE predicted high negative symptom scores across HC, ECP and CP groups, while high MCBN GE marginally predicted high negative symptoms across HC and CHR groups (Figure 1), indicating that these networks may differentially contribute to the development and maintenance of psychosis symptomatology^72^. We did not however find associations with positive symptoms across groups, contrary to our predictions and previous literature linking CB network dysfunction with positive symptom severity^28^. This suggests neurobiological markers for positive symptoms are constrained to changes within the CB-cortical networks and not necessarily related to CB-BG networks. Additional controls, such as number of relapses or time since diagnosis may show further relationships with symptomology we are not equipped to reveal.

We also investigated group differences in CCBN, MCBN, and cortical network GE. Main effects of group were found in both the CCBN and MCBN models (Figure 1, Table 2). CCBN GE was lower in CHR than ECP and marginally lower in CHR compared to HC. We cautiously posit this that deficit in CCBN GE in CHR is not related to age (which was controlled for) and may instead indicate risk of conversion that returns to normal levels after a first episode and conversion to ECP. However, the CHR sample did not differentiate between those who eventually converted so this argument needs further exploration with longitudinal follow-ups in larger samples. MCBN GE was marginally lower in chronic psychosis compared to HC and may be related to motor network dysfunction seen in patients with psychosis^72^. Cortical network (CON, DMN, EN, AN, and VN) GE showed a main effect of group but no group mean significantly differed f rom any other group mean (Figure 2, Table 2). These findings show psychosis-related brain network differences may be localized to CB-BG subcortical circuits and speak to the possible role these regions have in psychosis-related disorders and brain disease.

We were additionally interested in CB-BG GE as it related to cortical network efficiency across HC and the psychosis groups. We found associations between CCBN GE and CON, and AN GE (with a marginal association between CCBN GE and EN GE), an association between MCBN GE and AN GE, and a marginal association between the interaction of CCBN and MCBN GE and CON and AN GE (Figure 2, Table 2). Additionally, CCBN, MCBN, and their interaction each predicted VN GE within the ECP group (Figures 1 and 2, Tables 2 and 3). The relationships between and CCBN GE and CON, associated with attention, the EN, and the sensory networks (VN, AN) are supported by studies noting aberrations in sensory information processing, attention, and salience in psychotic disorders^73–76^. Two other cortical networks were also associated with CB-BG networks within the ECP group: the DMN and MN. The DMN exhibited an association with CCBN GE and marginal associations with MCBN GE and the interaction between CCBN and MCBN GE. MN GE was predicted by CCBN GE, and MCBN GE, and the interaction between CCBN and MCBN GE (Figure 2, Tables 2 and 3). Inefficient DMN suppression was related to schizophrenia symptom subtypes^77^, motor network deficiencies were seen in psychosis^31,72^, and deficient DMN suppression was noted in the CHR group while performing a learning task^46^. It may be that ECP brain networks are particularly variable and show differential effects within the CCBN, MCBN, DMN, and task-positive networks during the highly unstable time associated with first episodes and conversion to psychotic disorders that return to normal levels with treatment.

## Limitations

There are a few methodological limitations that should be addressed. First, combining three datasets comes with inherent difficulties. Although dataset membership was controlled for, differences in functional data collection parameters likely introduced noise. Relatedly, we worked to remove as much inter-dataset variance as possible by preprocessing the functional data using the same preprocessing steps. As the ADAPT dataset did not collect fieldmap images, no unwarping occurred. This is suboptimal and can be addressed in future studies by collecting data using the same parameters and by ensuring modern and best practices in image collection are followed. With that said, we believe the inclusion of this archival data that adds an additional psychosis spectrum group outweighed this limitation. Second, due to the structure of the combined dataset, there is a large correlation between dataset membership and group. For example, all those in the chronic psychosis group come from the COBRE dataset and all the CHR patients come from the ADAPT dataset. We accounted for this by controlling for dataset but this method may obscure additional group differences associated with disease state. Future studies may again need to use data from a single dataset or a larger number of datasets to decrease these correlations. Lastly, these data are not longitudinal so conclusions on disease progression within subjects cannot be assumed.

## Conclusion

In this study, we show that CB-BG GE predicts cognitive dysfunction and negative symptom scores. We also detailed relationships between CB-BG GE and cortical GE across groups and found both networks contribute to sensory, attentional, and motor networks, particularly within the ECP group. We propose CB-BG network GE may be a potential contributor to psychosis symptoms and further studies may determine it as a potential biomarker of a variety of psychiatric and neurological disorders.

## Supporting information

Supplemental Material 2: Stats code

## Acknowledgements and Funding Information

This work was supported in part by the Texas Virtual Data Library (ViDaL) funded by the Texas A&M University Research Development Fund; Data were provided [in part] by the Human Connectome Project, WU-Minn Consortium (Principal Investigators: David Van Essen and Kamil Ugurbil; 1U54MH091657) funded by the 16 NIH Institutes and Centers that support the NIH Blueprint for Neuroscience Research, and by the McDonnell Center for Systems Neuroscience at Washington University; Data was downloaded from the COllaborative Informatics and Neuroimaging Suite Data Exchange tool (COINS; http://coins.mrn.org/dx) and data collection was funded by R01MH084898-01A1; Further support for this work and the ADAPT data set came from R01MH094650, R33MH103231, R01MH120088 to V.A.M.; J.A.B. was supported in part by R01AG064010.

## Method

### Anatomical Data Preprocessing - fMRIPrep

A total of 1 T1-weighted (T1w) images were found within the input BIDS dataset. The T1-weighted (T1w) image was corrected for intensity non-uniformity (INU) with N4BiasFieldCorrection^1^, distributed with ANTs 2.3.3^2^ (RRID:SCR_004757), and used as T1w-reference throughout the workflow. The T1w-reference was then skull-stripped with a Nipype implementation of the antsBrainExtraction.sh workflow (from ANTs), using OASIS30ANTs as target template. Brain tissue segmentation of cerebrospinal fluid (CSF), white-matter (WM) and gray-matter (GM) was performed on the brain-extracted T1w using fast (FSL 6.0.5.1:57b01774, RRID:SCR_002823)^3^. Volume-based spatial normalization to two standard spaces (MNI152NLin6Asym, MNI152NLin2009cAsym) was performed through nonlinear registration with antsRegistration (ANTs 2.3.3), using brain-extracted versions of both T1w reference and the T1w template. The following templates were selected for spatial normalization: FSL’s MNI ICBM 152 non-linear 6th Generation Asymmetric Average Brain Stereotaxic Registration Model^4^, [RRID:SCR_002823; TemplateFlow ID: MNI152NLin6Asym], ICBM 152 Nonlinear Asymmetrical template version 2009c^5^ [RRID:SCR_008796; TemplateFlow ID: MNI152NLin2009cAsym].

### Functional Data Preprocessing - fMRIPrep

For each of the 1 BOLD runs found per subject (across all tasks and sessions), the following preprocessing was performed. First, a reference volume and its skull-stripped version were generated by aligning and averaging 1 single-band references (SBRefs). Head-motion parameters with respect to the BOLD reference (transformation matrices, and six corresponding rotation and translation parameters) are estimated before any spatiotemporal filtering using mcflirt (FSL 6.0.5.1:57b01774)^6^. BOLD runs were slice-time corrected to 0.346s (0.5 of slice acquisition range 0s-0.693s) using 3dTshift from AFNI^7^ (RRID:SCR_005927). The BOLD time-series (including slice-timing correction when applied) were resampled onto their original, native space by applying the transforms to correct for head-motion. These resampled BOLD time-series will be referred to as preprocessed BOLD in original space, or just preprocessed BOLD. The BOLD reference was then co-registered to the T1w reference using mri_coreg (FreeSurfer) followed by flirt (FSL 6.0.5.1:57b01774^8^ with the boundary-based registration^9^ cost-function. Co-registration was configured with six degrees of freedom. First, a reference volume and its skull-stripped version were generated using a custom methodology of fMRIPrep. Several confounding time-series were calculated based on the preprocessed BOLD: framewise displacement (FD), DVARS and three region-wise global signals. FD was computed using two formulations following Power (absolute sum of relative motions^10^) and Jenkinson (relative root mean square displacement between affines^6^. FD and DVARS are calculated for each functional run, both using their implementations in Nipype (following the definitions by^10^. The three global signals are extracted within the CSF, the WM, and the whole-brain masks. Additionally, a set of physiological regressors were extracted to allow for component-based noise correction (CompCor)^11^. Principal components are estimated after high-pass filtering the preprocessed BOLD time-series (using a discrete cosine filter with 128s cut-off) for the two CompCor variants: temporal (tCompCor) and anatomical (aCompCor). tCompCor components are then calculated from the top 2% variable voxels within the brain mask. For aCompCor, three probabilistic masks (CSF, WM and combined CSF+WM) are generated in anatomical space. The implementation differs from that of Behzadi et al. in that instead of eroding the masks by 2 pixels on BOLD space, the aCompCor masks are subtracted a mask of pixels that likely contain a volume fraction of GM. This mask is obtained by thresholding the corresponding partial volume map at 0.05, and it ensures components are not extracted from voxels containing a minimal fraction of GM. Finally, these masks are resampled into BOLD space and binarized by thresholding at 0.99 (as in the original implementation). Components are also calculated separately within the WM and CSF masks. For each CompCor decomposition, the k components with the largest singular values are retained, such that the retained components’ time series are sufficient to explain 50 percent of variance across the nuisance mask (CSF, WM, combined, or temporal). The remaining components are dropped from consideration. The head-motion estimates calculated in the correction step were also placed within the corresponding confounds file. The confound time series derived from head motion estimates and global signals were expanded with the inclusion of temporal derivatives and quadratic terms for each^12^. Frames that exceeded a threshold of 0.5 mm FD or 1.5 standardized DVARS were annotated as motion outliers. The BOLD time-series were resampled into standard space, generating a preprocessed BOLD run in MNI152NLin6Asym space. First, a reference volume and its skull-stripped version were generated using a custom methodology of fMRIPrep. All resamplings can be performed with a single interpolation step by composing all the pertinent transformations (i.e. head-motion transform matrices, susceptibility distortion correction when available, and co-registrations to anatomical and output spaces). Gridded (volumetric) resamplings were performed using antsApplyTransforms (ANTs), configured with Lanczos interpolation to minimize the smoothing effects of other kernels^13^. Non-gridded (surface) resamplings were performed using mri_vol2surf (FreeSurfer).

### CONN Preprocessing and Denoising

Additionally, structural images underwent tissue type segmentation and functional images underwent smoothing (5 mm FWHM) and artifact detection (global signal z-value threshold: 5, subject motion threshold: 0.9 mm) using the CONN toolbox (v. 21a; Whitfield-Gabrieli & Nieto-Castanon, 2012), a Matlab-based application designed for functional connectivity analysis. CONN was used with MATALB R2020a on Ubuntu 7.5. Data were then denoised using confound regressors for 5 temporal components each from the segmented CSF and white matter, 24 motion realignment parameters, signal and/or motion outliers, and the 1st order derivative from the effect of rest. Finally, data underwent linear detrending and bandpass filtering (0.008 - 0.09 Hz).

**Supplementary Table 1.**
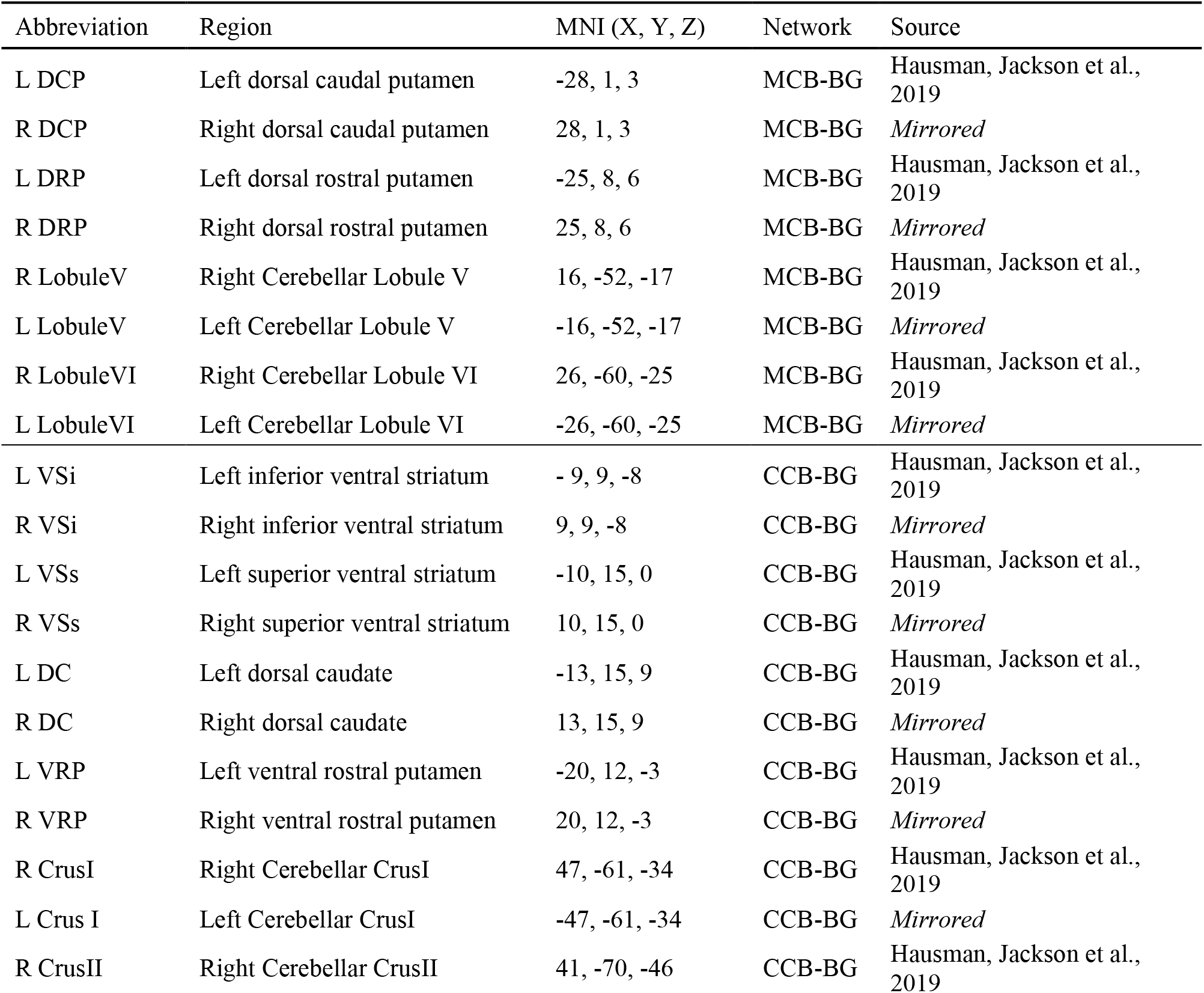
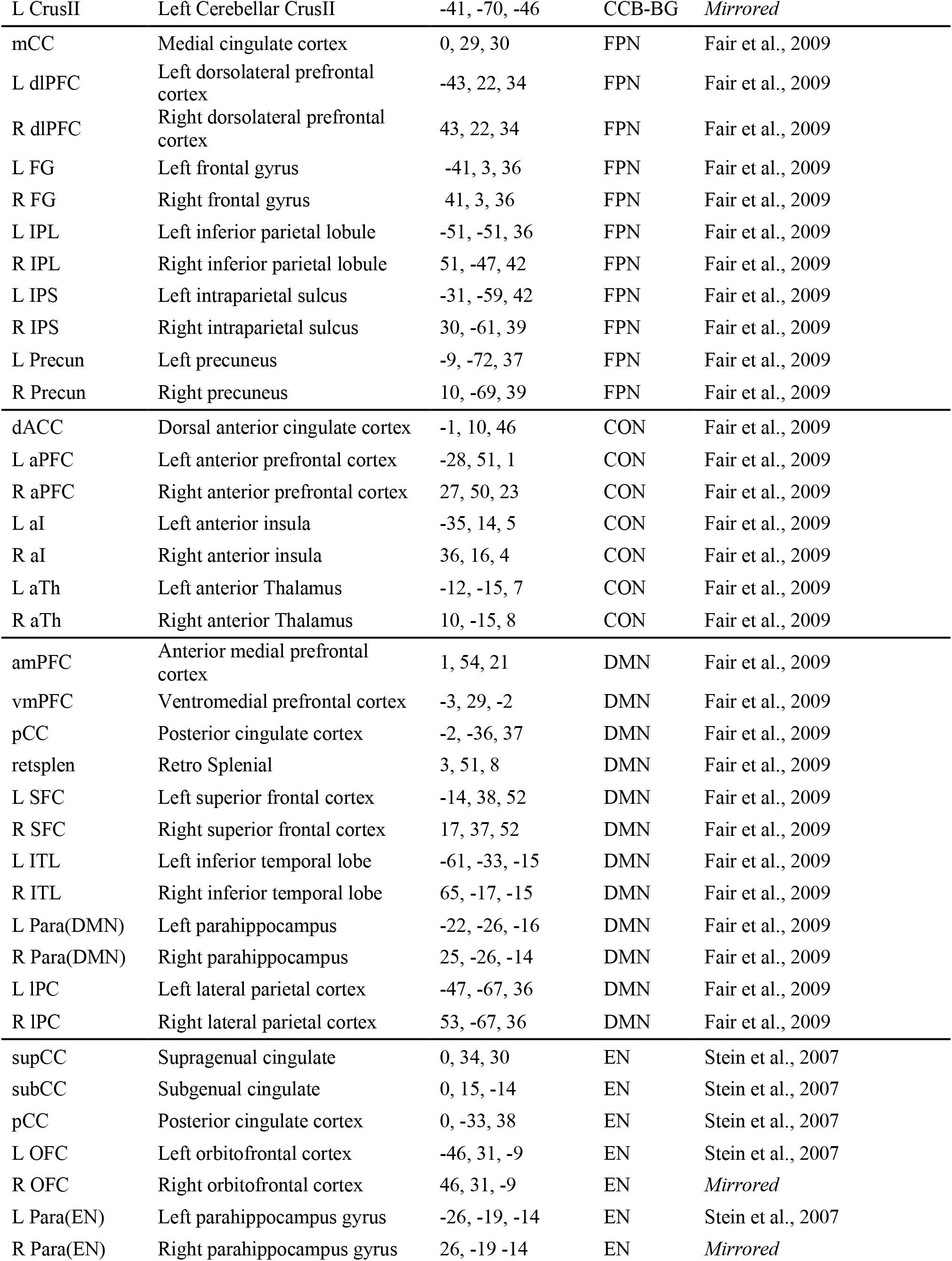
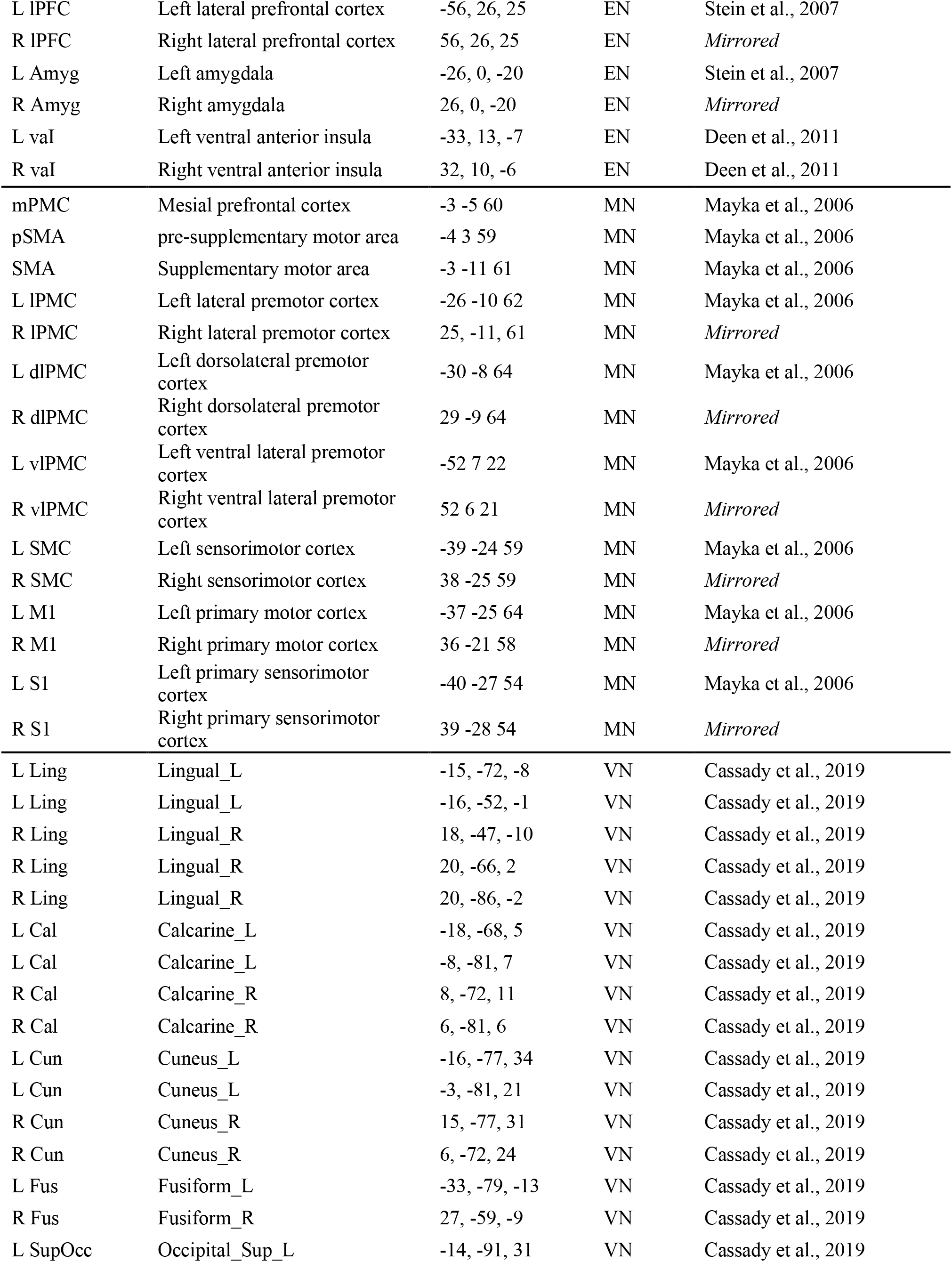
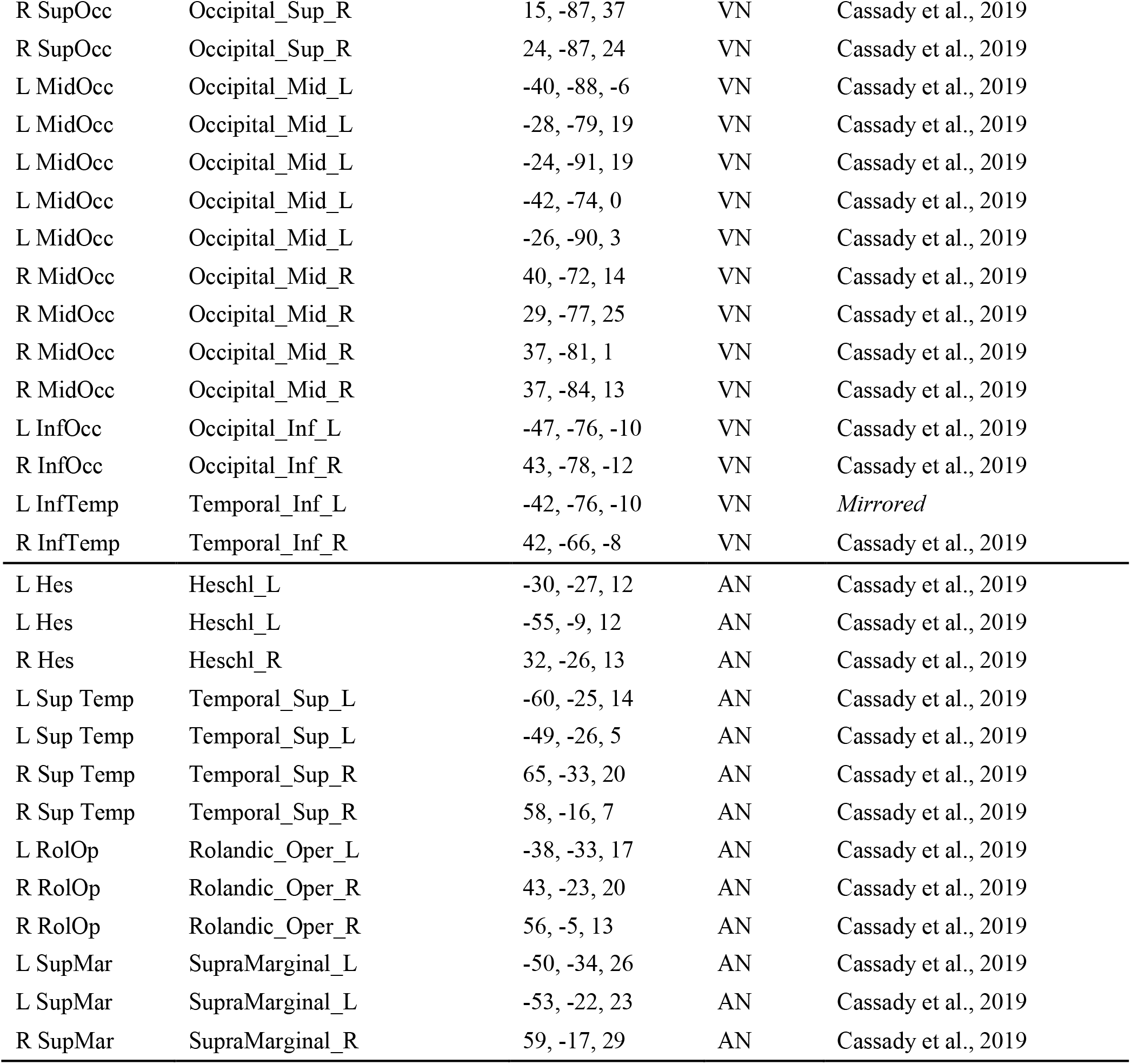
Abbreviations, region names, MNI coordinates, network association, and source for each ROIs used in the analyses. All ROIs are 3.5mm, spherical, and binarized. ROIs with the source “*Mirrored”* were created using its contralateral pair.

## Notes

**Compliance with Ethical Standards:** The authors declare no conflict of interest. The investigation reported here used data from human subjects that are publicly available.

### Competing Interest Statement

The authors have declared no competing interest.

