## Supplemental Material 2: Stats code for "Cerebello-Basal Ganglia Functional Network Integration in Psychosis"

Psychosis Project - Regression and ANOVA tables


### Psychosis Project - Regression and ANOVA tables

###### T. Bryan Jackson

### Load Libraries

```
library("ggplot2")
library("brms")
library("caret")
library("jtools")
library("multcomp")
library("emmeans")
library("apaTables")
library("papaja")
```

### Prepare Dataset

Set Up Group Factor, Split Dataset, etc.

#### Load Dataset

```
setwd("~/Desktop/PsychosisProject/Analyses")
raw_data <- read.csv("~/Desktop/PsychosisProject/Analyses/Data.csv")
dataset = raw_data
dataset$Group <- as.factor(dataset$Group)
dataset$Group <- factor(dataset$Group, levels=c('HC', 'CHR', 'ECP', 'CP'))

set.seed(4)

dataset_4 = split(dataset,dataset$Group)
CP = dataset_4$CP
ECP = dataset_4$ECP
CHR = dataset_4$CHR
HC = dataset_4$HC

noHC = rbind(CHR, ECP, CP)
```

#### Descriptive Stats

```
summary(dataset)
```

```
##      Subj             Dataset             Alcohol           Marj       
##  Length:448         Length:448         Min.   :0.000   Min.   :0.0000  
##  Class :character   Class :character   1st Qu.:0.000   1st Qu.:0.0000  
##  Mode  :character   Mode  :character   Median :1.000   Median :1.0000  
##                                        Mean   :0.877   Mean   :0.8993  
##                                        3rd Qu.:1.000   3rd Qu.:1.0000  
##                                        Max.   :3.000   Max.   :3.0000  
##                                        NA's   :1       NA's   :1       
##       Age            SIPS.P           SIPS.N          PANSS.P     
##  Min.   :12.00   Min.   : 0.000   Min.   : 0.000   Min.   : 0.00  
##  1st Qu.:19.00   1st Qu.: 0.000   1st Qu.: 0.000   1st Qu.: 0.00  
##  Median :22.00   Median : 5.000   Median : 2.000   Median :11.00  
##  Mean   :26.62   Mean   : 6.655   Mean   : 4.696   Mean   :10.22  
##  3rd Qu.:29.00   3rd Qu.:12.500   3rd Qu.: 8.000   3rd Qu.:16.00  
##  Max.   :65.00   Max.   :21.000   Max.   :23.000   Max.   :29.00  
##  NA's   :10      NA's   :277      NA's   :277      NA's   :172    
##     PANSS.N         PANSS.Cog    PANSS_Total_score      CCBN       
##  Min.   : 0.000   Min.   : 0.0   Min.   : 0.00     Min.   :0.3061  
##  1st Qu.: 0.000   1st Qu.: 9.0   1st Qu.:40.00     1st Qu.:0.6727  
##  Median :10.000   Median :10.0   Median :48.50     Median :0.7247  
##  Mean   : 9.616   Mean   :10.6   Mean   :46.98     Mean   :0.7151  
##  3rd Qu.:15.000   3rd Qu.:13.0   3rd Qu.:57.00     3rd Qu.:0.7803  
##  Max.   :29.000   Max.   :18.0   Max.   :78.00     Max.   :0.9470  
##  NA's   :172      NA's   :325    NA's   :326                       
##       MCBN             FPN              CON              DMN        
##  Min.   :0.1607   Min.   :0.3864   Min.   :0.2857   Min.   :0.3788  
##  1st Qu.:0.5893   1st Qu.:0.6481   1st Qu.:0.6429   1st Qu.:0.6812  
##  Median :0.7009   Median :0.7000   Median :0.7540   Median :0.7355  
##  Mean   :0.6760   Mean   :0.6952   Mean   :0.7287   Mean   :0.7315  
##  3rd Qu.:0.7812   3rd Qu.:0.7462   3rd Qu.:0.8333   3rd Qu.:0.7910  
##  Max.   :0.9821   Max.   :0.9364   Max.   :1.0000   Max.   :0.9545  
##                                                                     
##        EN               MN               VN               AN        
##  Min.   :0.2799   Min.   :0.4611   Min.   :0.6384   Min.   :0.4859  
##  1st Qu.:0.6067   1st Qu.:0.7298   1st Qu.:0.7660   1st Qu.:0.7158  
##  Median :0.6538   Median :0.7762   Median :0.8012   Median :0.7692  
##  Mean   :0.6447   Mean   :0.7769   Mean   :0.8047   Mean   :0.7654  
##  3rd Qu.:0.6923   3rd Qu.:0.8286   3rd Qu.:0.8449   3rd Qu.:0.8205  
##  Max.   :0.8205   Max.   :0.9810   Max.   :0.9536   Max.   :0.9615  
##                   NA's   :1                                         
##    CPZ_eqiuv           CBBG        Group    
##  Min.   :   0.0   Min.   :0.5031   HC :204  
##  1st Qu.:   0.0   1st Qu.:0.6652   CHR: 91  
##  Median :   0.0   Median :0.7000   ECP:104  
##  Mean   : 221.9   Mean   :0.6978   CP : 49  
##  3rd Qu.: 150.0   3rd Qu.:0.7336            
##  Max.   :7600.0   Max.   :0.8605            
##
```

### HC

```
summary(HC)
```

```
##      Subj             Dataset             Alcohol            Marj       
##  Length:204         Length:204         Min.   :0.0000   Min.   :0.0000  
##  Class :character   Class :character   1st Qu.:0.0000   1st Qu.:0.0000  
##  Mode  :character   Mode  :character   Median :1.0000   Median :1.0000  
##                                        Mean   :0.7931   Mean   :0.8079  
##                                        3rd Qu.:1.0000   3rd Qu.:1.0000  
##                                        Max.   :3.0000   Max.   :3.0000  
##                                        NA's   :1        NA's   :1       
##       Age            SIPS.P         SIPS.N          PANSS.P      
##  Min.   :12.00   Min.   :0.00   Min.   :0.0000   Min.   : 0.000  
##  1st Qu.:19.00   1st Qu.:0.00   1st Qu.:0.0000   1st Qu.: 0.000  
##  Median :22.38   Median :0.00   Median :0.0000   Median : 0.000  
##  Mean   :27.72   Mean   :0.60   Mean   :0.4375   Mean   : 4.163  
##  3rd Qu.:34.25   3rd Qu.:0.25   3rd Qu.:1.0000   3rd Qu.: 9.000  
##  Max.   :65.00   Max.   :5.00   Max.   :3.0000   Max.   :25.000  
##  NA's   :8       NA's   :124    NA's   :124      NA's   :81      
##     PANSS.N         PANSS.Cog     PANSS_Total_score      CCBN       
##  Min.   : 0.000   Min.   : 0.00   Min.   : 0.00     Min.   :0.3061  
##  1st Qu.: 0.000   1st Qu.: 9.00   1st Qu.:36.00     1st Qu.:0.6837  
##  Median : 0.000   Median : 9.00   Median :46.00     Median :0.7361  
##  Mean   : 3.675   Mean   : 9.07   Mean   :41.53     Mean   :0.7266  
##  3rd Qu.: 7.000   3rd Qu.:10.00   3rd Qu.:53.50     3rd Qu.:0.7854  
##  Max.   :23.000   Max.   :16.00   Max.   :78.00     Max.   :0.9015  
##  NA's   :81       NA's   :161     NA's   :161                       
##       MCBN             FPN              CON              DMN        
##  Min.   :0.1607   Min.   :0.3864   Min.   :0.2857   Min.   :0.5013  
##  1st Qu.:0.6119   1st Qu.:0.6538   1st Qu.:0.6667   1st Qu.:0.6856  
##  Median :0.7143   Median :0.7030   Median :0.7778   Median :0.7538  
##  Mean   :0.6915   Mean   :0.6971   Mean   :0.7473   Mean   :0.7398  
##  3rd Qu.:0.8036   3rd Qu.:0.7485   3rd Qu.:0.8512   3rd Qu.:0.8005  
##  Max.   :0.9821   Max.   :0.9000   Max.   :1.0000   Max.   :0.9545  
##                                                                     
##        EN               MN               VN               AN        
##  Min.   :0.4509   Min.   :0.5183   Min.   :0.6771   Min.   :0.5365  
##  1st Qu.:0.6152   1st Qu.:0.7355   1st Qu.:0.7681   1st Qu.:0.7265  
##  Median :0.6608   Median :0.7825   Median :0.8034   Median :0.7842  
##  Mean   :0.6550   Mean   :0.7809   Mean   :0.8061   Mean   :0.7795  
##  3rd Qu.:0.7014   3rd Qu.:0.8230   3rd Qu.:0.8435   3rd Qu.:0.8280  
##  Max.   :0.7756   Max.   :0.9810   Max.   :0.9526   Max.   :0.9615  
##                                                                     
##    CPZ_eqiuv      CBBG        Group    
##  Min.   :0   Min.   :0.5031   HC :204  
##  1st Qu.:0   1st Qu.:0.6711   CHR:  0  
##  Median :0   Median :0.7096   ECP:  0  
##  Mean   :0   Mean   :0.7066   CP :  0  
##  3rd Qu.:0   3rd Qu.:0.7400            
##  Max.   :0   Max.   :0.8605            
##
```

```
sd(HC$Age, na.rm=TRUE)
```

```
## [1] 12.4743
```

```
sd(HC$Alcohol, na.rm=TRUE)
```

```
## [1] 0.7938824
```

```
sd(HC$Marj, na.rm=TRUE)
```

```
## [1] 0.8660465
```

```
sd(HC$CPZ_eqiuv, na.rm=TRUE)
```

```
## [1] 0
```

```
sd(HC$PANSS.P, na.rm=TRUE)
```

```
## [1] 6.420222
```

```
sd(HC$PANSS.N, na.rm=TRUE)
```

```
## [1] 5.940271
```

```
sd(HC$PANSS.Cog, na.rm=TRUE)
```

```
## [1] 2.676132
```

```
sd(HC$SIPS.P, na.rm=TRUE)
```

```
## [1] 1.268908
```

```
sd(HC$SIPS.N, na.rm=TRUE)
```

```
## [1] 0.7604754
```

##### CHR

```
summary(CHR)
```

```
##      Subj             Dataset             Alcohol            Marj       
##  Length:91          Length:91          Min.   :0.0000   Min.   :0.0000  
##  Class :character   Class :character   1st Qu.:0.0000   1st Qu.:0.0000  
##  Mode  :character   Mode  :character   Median :0.0000   Median :0.0000  
##                                        Mean   :0.3956   Mean   :0.1319  
##                                        3rd Qu.:1.0000   3rd Qu.:0.0000  
##                                        Max.   :1.0000   Max.   :1.0000  
##                                                                         
##       Age            SIPS.P          SIPS.N         PANSS.P       PANSS.N   
##  Min.   :13.00   Min.   : 0.00   Min.   : 0.00   Min.   : NA   Min.   : NA  
##  1st Qu.:18.50   1st Qu.: 9.00   1st Qu.: 3.00   1st Qu.: NA   1st Qu.: NA  
##  Median :19.00   Median :12.00   Median : 7.00   Median : NA   Median : NA  
##  Mean   :19.51   Mean   :11.98   Mean   : 8.44   Mean   :NaN   Mean   :NaN  
##  3rd Qu.:21.00   3rd Qu.:15.00   3rd Qu.:13.00   3rd Qu.: NA   3rd Qu.: NA  
##  Max.   :24.00   Max.   :21.00   Max.   :23.00   Max.   : NA   Max.   : NA  
##                                                  NA's   :91    NA's   :91   
##    PANSS.Cog   PANSS_Total_score      CCBN             MCBN       
##  Min.   : NA   Min.   : NA       Min.   :0.3561   Min.   :0.2649  
##  1st Qu.: NA   1st Qu.: NA       1st Qu.:0.6290   1st Qu.:0.5920  
##  Median : NA   Median : NA       Median :0.7146   Median :0.7024  
##  Mean   :NaN   Mean   :NaN       Mean   :0.6936   Mean   :0.6886  
##  3rd Qu.: NA   3rd Qu.: NA       3rd Qu.:0.7790   3rd Qu.:0.8274  
##  Max.   : NA   Max.   : NA       Max.   :0.8636   Max.   :0.9821  
##  NA's   :91    NA's   :91                                         
##       FPN              CON              DMN               EN        
##  Min.   :0.3924   Min.   :0.2857   Min.   :0.4545   Min.   :0.2799  
##  1st Qu.:0.6571   1st Qu.:0.6290   1st Qu.:0.6982   1st Qu.:0.5790  
##  Median :0.7000   Median :0.7222   Median :0.7475   Median :0.6357  
##  Mean   :0.7061   Mean   :0.7011   Mean   :0.7456   Mean   :0.6216  
##  3rd Qu.:0.7576   3rd Qu.:0.8214   3rd Qu.:0.7942   3rd Qu.:0.6784  
##  Max.   :0.8818   Max.   :1.0000   Max.   :0.9167   Max.   :0.7821  
##                                                                     
##        MN               VN               AN           CPZ_eqiuv
##  Min.   :0.6294   Min.   :0.6781   Min.   :0.4861   Min.   :0  
##  1st Qu.:0.7194   1st Qu.:0.7636   1st Qu.:0.7131   1st Qu.:0  
##  Median :0.7786   Median :0.8155   Median :0.7735   Median :0  
##  Mean   :0.7807   Mean   :0.8137   Mean   :0.7641   Mean   :0  
##  3rd Qu.:0.8333   3rd Qu.:0.8668   3rd Qu.:0.8205   3rd Qu.:0  
##  Max.   :0.9619   Max.   :0.9536   Max.   :0.9359   Max.   :0  
##  NA's   :1                                                     
##       CBBG        Group   
##  Min.   :0.5281   HC : 0  
##  1st Qu.:0.6442   CHR:91  
##  Median :0.6886   ECP: 0  
##  Mean   :0.6815   CP : 0  
##  3rd Qu.:0.7274           
##  Max.   :0.8316           
##
```

```
sd(CHR$Age, na.rm=TRUE)
```

```
## [1] 2.115412
```

```
sd(CHR$Alcohol, na.rm=TRUE)
```

```
## [1] 0.4916892
```

```
sd(CHR$Marj, na.rm=TRUE)
```

```
## [1] 0.3402219
```

```
sd(CHR$CPZ_eqiuv, na.rm=TRUE)
```

```
## [1] 0
```

```
sd(CHR$PANSS.P, na.rm=TRUE)
```

```
## [1] NA
```

```
sd(CHR$PANSS.N, na.rm=TRUE)
```

```
## [1] NA
```

```
sd(CHR$PANSS.Cog, na.rm=TRUE)
```

```
## [1] NA
```

```
sd(CHR$SIPS.P, na.rm=TRUE)
```

```
## [1] 4.464621
```

```
sd(CHR$SIPS.N, na.rm=TRUE)
```

```
## [1] 6.347718
```

##### ECP

```
summary(ECP)
```

```
##      Subj             Dataset             Alcohol           Marj      
##  Length:104         Length:104         Min.   :1.000   Min.   :1.000  
##  Class :character   Class :character   1st Qu.:1.000   1st Qu.:1.000  
##  Mode  :character   Mode  :character   Median :1.000   Median :1.000  
##                                        Mean   :1.231   Mean   :1.529  
##                                        3rd Qu.:1.000   3rd Qu.:2.000  
##                                        Max.   :3.000   Max.   :3.000  
##                                                                       
##       Age            SIPS.P        SIPS.N       PANSS.P         PANSS.N     
##  Min.   :17.92   Min.   : NA   Min.   : NA   Min.   : 0.00   Min.   : 0.00  
##  1st Qu.:20.10   1st Qu.: NA   1st Qu.: NA   1st Qu.:12.00   1st Qu.: 9.00  
##  Median :22.00   Median : NA   Median : NA   Median :15.00   Median :14.00  
##  Mean   :22.71   Mean   :NaN   Mean   :NaN   Mean   :15.07   Mean   :14.11  
##  3rd Qu.:24.17   3rd Qu.: NA   3rd Qu.: NA   3rd Qu.:18.25   3rd Qu.:18.00  
##  Max.   :34.08   Max.   : NA   Max.   : NA   Max.   :29.00   Max.   :29.00  
##  NA's   :2       NA's   :104   NA's   :104                                  
##    PANSS.Cog     PANSS_Total_score      CCBN             MCBN       
##  Min.   : 0.00   Min.   : 0.00     Min.   :0.4235   Min.   :0.2292  
##  1st Qu.: 9.00   1st Qu.:42.00     1st Qu.:0.6739   1st Qu.:0.5482  
##  Median :11.00   Median :51.00     Median :0.7121   Median :0.6637  
##  Mean   :11.43   Mean   :49.95     Mean   :0.7138   Mean   :0.6340  
##  3rd Qu.:14.00   3rd Qu.:60.00     3rd Qu.:0.7626   3rd Qu.:0.7359  
##  Max.   :18.00   Max.   :77.00     Max.   :0.9470   Max.   :0.9821  
##  NA's   :24      NA's   :25                                         
##       FPN              CON              DMN               EN        
##  Min.   :0.4152   Min.   :0.2857   Min.   :0.3788   Min.   :0.3440  
##  1st Qu.:0.6273   1st Qu.:0.5815   1st Qu.:0.6509   1st Qu.:0.6035  
##  Median :0.6902   Median :0.7381   Median :0.7172   Median :0.6407  
##  Mean   :0.6825   Mean   :0.7016   Mean   :0.7052   Mean   :0.6373  
##  3rd Qu.:0.7424   3rd Qu.:0.8115   3rd Qu.:0.7626   3rd Qu.:0.6830  
##  Max.   :0.9364   Max.   :1.0000   Max.   :0.9545   Max.   :0.8205  
##                                                                     
##        MN               VN               AN           CPZ_eqiuv     
##  Min.   :0.4611   Min.   :0.6384   Min.   :0.4859   Min.   :   0.0  
##  1st Qu.:0.7079   1st Qu.:0.7704   1st Qu.:0.6856   1st Qu.:   0.0  
##  Median :0.7579   Median :0.7982   Median :0.7489   Median : 300.0  
##  Mean   :0.7629   Mean   :0.8047   Mean   :0.7467   Mean   : 411.2  
##  3rd Qu.:0.8321   3rd Qu.:0.8343   3rd Qu.:0.8077   3rd Qu.: 400.0  
##  Max.   :0.9286   Max.   :0.9315   Max.   :0.9359   Max.   :4797.0  
##                                                                     
##       CBBG        Group    
##  Min.   :0.5219   HC :  0  
##  1st Qu.:0.6592   CHR:  0  
##  Median :0.6895   ECP:104  
##  Mean   :0.6874   CP :  0  
##  3rd Qu.:0.7195            
##  Max.   :0.8228            
##
```

```
sd(ECP$Age, na.rm=TRUE)
```

```
## [1] 3.438597
```

```
sd(ECP$Alcohol, na.rm=TRUE)
```

```
## [1] 0.6267619
```

```
sd(ECP$Marj, na.rm=TRUE)
```

```
## [1] 0.858501
```

```
sd(ECP$CPZ_eqiuv, na.rm=TRUE)
```

```
## [1] 648.8351
```

```
sd(ECP$PANSS.P, na.rm=TRUE)
```

```
## [1] 5.120423
```

```
sd(ECP$PANSS.N, na.rm=TRUE)
```

```
## [1] 5.923326
```

```
sd(ECP$PANSS.Cog, na.rm=TRUE)
```

```
## [1] 3.279298
```

```
sd(ECP$SIPS.P, na.rm=TRUE)
```

```
## [1] NA
```

```
sd(ECP$SIPS.N, na.rm=TRUE)
```

```
## [1] NA
```

### CP

```
summary(CP)
```

```
##      Subj             Dataset             Alcohol           Marj      
##  Length:49          Length:49          Min.   :1.000   Min.   :1.000  
##  Class :character   Class :character   1st Qu.:1.000   1st Qu.:1.000  
##  Mode  :character   Mode  :character   Median :1.000   Median :1.000  
##                                        Mean   :1.367   Mean   :1.367  
##                                        3rd Qu.:2.000   3rd Qu.:1.000  
##                                        Max.   :3.000   Max.   :3.000  
##                                                                       
##       Age            SIPS.P        SIPS.N       PANSS.P         PANSS.N  
##  Min.   :19.00   Min.   : NA   Min.   : NA   Min.   : 7.00   Min.   : 7  
##  1st Qu.:36.00   1st Qu.: NA   1st Qu.: NA   1st Qu.:13.00   1st Qu.:11  
##  Median :44.00   Median : NA   Median : NA   Median :15.00   Median :14  
##  Mean   :43.57   Mean   :NaN   Mean   :NaN   Mean   :15.14   Mean   :15  
##  3rd Qu.:52.00   3rd Qu.: NA   3rd Qu.: NA   3rd Qu.:18.00   3rd Qu.:18  
##  Max.   :64.00   Max.   : NA   Max.   : NA   Max.   :25.00   Max.   :29  
##                  NA's   :49    NA's   :49                                
##    PANSS.Cog   PANSS_Total_score      CCBN             MCBN       
##  Min.   : NA   Min.   : NA       Min.   :0.4545   Min.   :0.3036  
##  1st Qu.: NA   1st Qu.: NA       1st Qu.:0.6591   1st Qu.:0.6381  
##  Median : NA   Median : NA       Median :0.7273   Median :0.7143  
##  Mean   :NaN   Mean   :NaN       Mean   :0.7101   Mean   :0.6769  
##  3rd Qu.: NA   3rd Qu.: NA       3rd Qu.:0.7727   3rd Qu.:0.7738  
##  Max.   : NA   Max.   : NA       Max.   :0.8409   Max.   :0.9643  
##  NA's   :49    NA's   :49                                         
##       FPN              CON              DMN               EN        
##  Min.   :0.4576   Min.   :0.3651   Min.   :0.5066   Min.   :0.4676  
##  1st Qu.:0.6394   1st Qu.:0.6905   1st Qu.:0.6894   1st Qu.:0.6346  
##  Median :0.7000   Median :0.7778   Median :0.7285   Median :0.6667  
##  Mean   :0.6947   Mean   :0.7603   Mean   :0.7264   Mean   :0.6605  
##  3rd Qu.:0.7364   3rd Qu.:0.8492   3rd Qu.:0.7727   3rd Qu.:0.6902  
##  Max.   :0.8545   Max.   :1.0000   Max.   :0.8636   Max.   :0.7714  
##                                                                     
##        MN               VN               AN           CPZ_eqiuv   
##  Min.   :0.6484   Min.   :0.6808   Min.   :0.5107   Min.   :   0  
##  1st Qu.:0.7571   1st Qu.:0.7446   1st Qu.:0.7179   1st Qu.: 268  
##  Median :0.7794   Median :0.7776   Median :0.7564   Median : 600  
##  Mean   :0.7830   Mean   :0.7819   Mean   :0.7488   Mean   :1156  
##  3rd Qu.:0.8286   3rd Qu.:0.8246   3rd Qu.:0.7863   3rd Qu.:1600  
##  Max.   :0.8857   Max.   :0.9224   Max.   :0.9231   Max.   :7600  
##                                                                   
##       CBBG        Group   
##  Min.   :0.5426   HC : 0  
##  1st Qu.:0.6851   CHR: 0  
##  Median :0.7096   ECP: 0  
##  Mean   :0.7133   CP :49  
##  3rd Qu.:0.7430           
##  Max.   :0.7921           
##
```

```
sd(CP$Age, na.rm=TRUE)
```

```
## [1] 10.45028
```

```
sd(CP$Alcohol, na.rm=TRUE)
```

```
## [1] 0.6675165
```

```
sd(CP$Marj, na.rm=TRUE)
```

```
## [1] 0.6980293
```

```
sd(CP$CPZ_eqiuv, na.rm=TRUE)
```

```
## [1] 1399.323
```

```
sd(CP$PANSS.P, na.rm=TRUE)
```

```
## [1] 4.067145
```

```
sd(CP$PANSS.N, na.rm=TRUE)
```

```
## [1] 5.303301
```

```
sd(CP$PANSS.Cog, na.rm=TRUE)
```

```
## [1] NA
```

```
sd(CP$SIPS.P, na.rm=TRUE)
```

```
## [1] NA
```

```
sd(CP$SIPS.N, na.rm=TRUE)
```

```
## [1] NA
```

### Group Differences in Symptoms

#### PANSS-P

```
GroupDiffPP <- aov(PANSS.P ~ Group + Alcohol+ Marj + Age + CPZ_eqiuv, dataset)
summary(GroupDiffPP)
```

```
##              Df Sum Sq Mean Sq F value   Pr(>F)    
## Group         2   8589    4294 174.617  < 2e-16 ***
## Alcohol       1     49      49   1.987 0.159820    
## Marj          1    326     326  13.239 0.000331 ***
## Age           1    832     832  33.847 1.75e-08 ***
## CPZ_eqiuv     1      2       2   0.072 0.788757    
## Residuals   260   6394      25                     
## ---
## Signif. codes:  0 '***' 0.001 '**' 0.01 '*' 0.05 '.' 0.1 ' ' 1
## 181 observations deleted due to missingness
```

```
PANSS_P_group_plot <- effect_plot(GroupDiffPP, pred = Group, interval = TRUE, partial.residuals = TRUE, jitter = .2)
PANSS_P_group_plot + theme(text = element_text(size = 24), panel.border = element_rect(colour = "black", fill=NA, size=1), axis.title.x = element_blank(), axis.title.y = element_blank())
```

```
posthoc_PP <- glht(GroupDiffPP, linfct = mcp(Group="Tukey"))
summary(posthoc_PP, adjusted(type='fdr'))
```

```
## 
##   Simultaneous Tests for General Linear Hypotheses
## 
## Multiple Comparisons of Means: Tukey Contrasts
## 
## 
## Fit: aov(formula = PANSS.P ~ Group + Alcohol + Marj + Age + CPZ_eqiuv, 
##     data = dataset)
## 
## Linear Hypotheses:
##               Estimate Std. Error t value Pr(>|t|)    
## ECP - HC == 0    9.137      0.782  11.684  < 2e-16 ***
## CP - HC == 0    12.990      1.026  12.657  < 2e-16 ***
## CP - ECP == 0    3.853      1.133   3.401 0.000776 ***
## ---
## Signif. codes:  0 '***' 0.001 '**' 0.01 '*' 0.05 '.' 0.1 ' ' 1
## (Adjusted p values reported -- fdr method)
```

#### PANSS-N

```
GroupDiffPN <- aov(PANSS.N ~ Group + Alcohol+ Marj + Age + CPZ_eqiuv, dataset)
summary(GroupDiffPN)
```

```
##              Df Sum Sq Mean Sq F value   Pr(>F)    
## Group         2   7852    3926 130.054  < 2e-16 ***
## Alcohol       1      2       2   0.065  0.79929    
## Marj          1    252     252   8.345  0.00419 ** 
## Age           1    864     864  28.620 1.93e-07 ***
## CPZ_eqiuv     1      4       4   0.125  0.72426    
## Residuals   260   7849      30                     
## ---
## Signif. codes:  0 '***' 0.001 '**' 0.01 '*' 0.05 '.' 0.1 ' ' 1
## 181 observations deleted due to missingness
```

```
PANSS_N_group_plot <- effect_plot(GroupDiffPN, pred = Group, interval = TRUE, partial.residuals = TRUE, jitter = .2)
PANSS_N_group_plot + theme(text = element_text(size = 24), panel.border = element_rect(colour = "black", fill=NA, size=1), axis.title.x = element_blank(), axis.title.y = element_blank())
```

```
posthoc_PN <- glht(GroupDiffPN, linfct = mcp(Group="Tukey"))
summary(posthoc_PN, adjusted(type='fdr'))
```

```
## 
##   Simultaneous Tests for General Linear Hypotheses
## 
## Multiple Comparisons of Means: Tukey Contrasts
## 
## 
## Fit: aov(formula = PANSS.N ~ Group + Alcohol + Marj + Age + CPZ_eqiuv, 
##     data = dataset)
## 
## Linear Hypotheses:
##               Estimate Std. Error t value Pr(>|t|)    
## ECP - HC == 0   8.2279     0.8664   9.497  < 2e-16 ***
## CP - HC == 0   13.1886     1.1370  11.600  < 2e-16 ***
## CP - ECP == 0   4.9607     1.2551   3.953 9.97e-05 ***
## ---
## Signif. codes:  0 '***' 0.001 '**' 0.01 '*' 0.05 '.' 0.1 ' ' 1
## (Adjusted p values reported -- fdr method)
```

#### PANSS-Cog

```
GroupDiffPC <- aov(PANSS.Cog ~ Group + Alcohol+ Marj + Age  + CPZ_eqiuv, dataset)
summary(GroupDiffPC)
```

```
##              Df Sum Sq Mean Sq F value   Pr(>F)    
## Group         1  113.4  113.36  14.763 0.000206 ***
## Alcohol       1    0.1    0.06   0.008 0.928996    
## Marj          1    0.6    0.56   0.072 0.788248    
## Age           1    4.4    4.38   0.571 0.451695    
## CPZ_eqiuv     1   83.4   83.44  10.866 0.001326 ** 
## Residuals   108  829.3    7.68                     
## ---
## Signif. codes:  0 '***' 0.001 '**' 0.01 '*' 0.05 '.' 0.1 ' ' 1
## 334 observations deleted due to missingness
```

```
PANSS_Cog_group_plot <- effect_plot(GroupDiffPC, pred = Group, interval = TRUE, partial.residuals = TRUE, jitter = .2)
PANSS_Cog_group_plot + theme(text = element_text(size = 24), panel.border = element_rect(colour = "black", fill=NA, size=1), axis.title.x = element_blank(), axis.title.y = element_blank())
```

#### SIPS-P

```
GroupDiffSP <- aov(SIPS.P ~ Group + Alcohol + Marj + Age , dataset)
summary(GroupDiffSP)
```

```
##              Df Sum Sq Mean Sq F value  Pr(>F)    
## Group         1   5511    5511 515.689 < 2e-16 ***
## Alcohol       1      3       3   0.306 0.58101    
## Marj          1    108     108  10.110 0.00176 ** 
## Age           1     36      36   3.340 0.06943 .  
## Residuals   166   1774      11                    
## ---
## Signif. codes:  0 '***' 0.001 '**' 0.01 '*' 0.05 '.' 0.1 ' ' 1
## 277 observations deleted due to missingness
```

```
SIPS_P_group_plot <- effect_plot(GroupDiffSP, pred = Group, interval = TRUE, partial.residuals = TRUE, jitter = .2)
SIPS_P_group_plot + theme(text = element_text(size = 24), panel.border = element_rect(colour = "black", fill=NA, size=1), axis.title.x = element_blank(), axis.title.y = element_blank())
```

```
posthoc_SP <- glht(GroupDiffSP, linfct = mcp(Group="Tukey"))
summary(posthoc_SP, adjusted(type='fdr'))
```

```
## 
##   Simultaneous Tests for General Linear Hypotheses
## 
## Multiple Comparisons of Means: Tukey Contrasts
## 
## 
## Fit: aov(formula = SIPS.P ~ Group + Alcohol + Marj + Age, data = dataset)
## 
## Linear Hypotheses:
##               Estimate Std. Error t value Pr(>|t|)    
## CHR - HC == 0  10.6776     0.5599   19.07   <2e-16 ***
## ---
## Signif. codes:  0 '***' 0.001 '**' 0.01 '*' 0.05 '.' 0.1 ' ' 1
## (Adjusted p values reported -- fdr method)
```

#### SIPS-N

```
GroupDiffSN <- aov(SIPS.N ~ Group + Alcohol+ Marj + Age, dataset)
summary(GroupDiffSN)
```

```
##              Df Sum Sq Mean Sq F value  Pr(>F)    
## Group         1   2726  2726.1 131.540 < 2e-16 ***
## Alcohol       1     32    32.2   1.555 0.21421    
## Marj          1    199   199.3   9.618 0.00226 ** 
## Age           1      0     0.3   0.016 0.90054    
## Residuals   166   3440    20.7                    
## ---
## Signif. codes:  0 '***' 0.001 '**' 0.01 '*' 0.05 '.' 0.1 ' ' 1
## 277 observations deleted due to missingness
```

```
SIPS_N_group_plot <- effect_plot(GroupDiffSN, pred = Group, interval = TRUE, partial.residuals = TRUE, jitter = .2)
SIPS_N_group_plot + theme(text = element_text(size = 24), panel.border = element_rect(colour = "black", fill=NA, size=1), axis.title.x = element_blank(), axis.title.y = element_blank())
```

```
posthoc_SN <- glht(GroupDiffSN, linfct = mcp(Group="Tukey"))
summary(posthoc_SN, adjusted(type='fdr'))
```

```
## 
##   Simultaneous Tests for General Linear Hypotheses
## 
## Multiple Comparisons of Means: Tukey Contrasts
## 
## 
## Fit: aov(formula = SIPS.N ~ Group + Alcohol + Marj + Age, data = dataset)
## 
## Linear Hypotheses:
##               Estimate Std. Error t value Pr(>|t|)    
## CHR - HC == 0   7.0730     0.7796   9.072 4.44e-16 ***
## ---
## Signif. codes:  0 '***' 0.001 '**' 0.01 '*' 0.05 '.' 0.1 ' ' 1
## (Adjusted p values reported -- fdr method)
```

### Group Differences in CB-BG GE

#### CCBN

```
GroupDiffC <- aov(CCBN ~ Group + Alcohol+ Marj + Age + Dataset + CPZ_eqiuv, dataset)
summary(GroupDiffC)
```

```
##              Df Sum Sq Mean Sq F value   Pr(>F)    
## Group         3  0.072 0.02416   2.913 0.034138 *  
## Alcohol       1  0.006 0.00569   0.686 0.407850    
## Marj          1  0.042 0.04161   5.016 0.025622 *  
## Age           1  0.009 0.00948   1.143 0.285676    
## Dataset       2  0.142 0.07093   8.552 0.000228 ***
## CPZ_eqiuv     1  0.010 0.00988   1.191 0.275804    
## Residuals   428  3.550 0.00829                     
## ---
## Signif. codes:  0 '***' 0.001 '**' 0.01 '*' 0.05 '.' 0.1 ' ' 1
## 10 observations deleted due to missingness
```

```
CCBN_group_plot <- effect_plot(GroupDiffC, pred = Group, interval = TRUE, partial.residuals = TRUE, jitter = .2)
CCBN_group_plot + ylab("CCBN GE") + theme(text = element_text(size = 24), panel.border = element_rect(colour = "black", fill=NA, size=1), axis.title.x = element_blank())
```

```
posthoc_C <- glht(GroupDiffC, linfct = mcp(Group="Tukey"))
summary(posthoc_C, adjusted(type='fdr'))
```

```
## 
##   Simultaneous Tests for General Linear Hypotheses
## 
## Multiple Comparisons of Means: Tukey Contrasts
## 
## 
## Fit: aov(formula = CCBN ~ Group + Alcohol + Marj + Age + Dataset + 
##     CPZ_eqiuv, data = dataset)
## 
## Linear Hypotheses:
##                Estimate Std. Error t value Pr(>|t|)  
## CHR - HC == 0  -0.03594    0.01417  -2.537   0.0346 *
## ECP - HC == 0   0.01993    0.01511   1.318   0.2821  
## CP - HC == 0   -0.01912    0.01882  -1.016   0.3724  
## ECP - CHR == 0  0.05587    0.02090   2.673   0.0346 *
## CP - CHR == 0   0.01682    0.02342   0.718   0.4730  
## CP - ECP == 0  -0.03905    0.02143  -1.822   0.1382  
## ---
## Signif. codes:  0 '***' 0.001 '**' 0.01 '*' 0.05 '.' 0.1 ' ' 1
## (Adjusted p values reported -- fdr method)
```

#### MCBN

```
GroupDiffM <- aov(MCBN ~ Group + Alcohol+ Marj + Age + Dataset + CPZ_eqiuv, dataset)
summary(GroupDiffM)
```

```
##              Df Sum Sq Mean Sq F value   Pr(>F)    
## Group         3  0.257 0.08575   3.972  0.00823 ** 
## Alcohol       1  0.002 0.00200   0.093  0.76112    
## Marj          1  0.052 0.05180   2.399  0.12214    
## Age           1  0.078 0.07830   3.626  0.05755 .  
## Dataset       2  0.531 0.26529  12.287 6.48e-06 ***
## CPZ_eqiuv     1  0.050 0.05038   2.333  0.12737    
## Residuals   428  9.241 0.02159                     
## ---
## Signif. codes:  0 '***' 0.001 '**' 0.01 '*' 0.05 '.' 0.1 ' ' 1
## 10 observations deleted due to missingness
```

```
MCBN_group_plot <- effect_plot(GroupDiffM, pred = Group, interval = TRUE, partial.residuals = TRUE, jitter = .2)
MCBN_group_plot + ylab("MCBN GE") + theme(text = element_text(size = 24), panel.border = element_rect(colour = "black", fill=NA, size=1), axis.title.x = element_blank())
```

```
posthoc_M <- glht(GroupDiffM, linfct = mcp(Group="Tukey"))
summary(posthoc_M, adjusted(type='fdr'))
```

```
## 
##   Simultaneous Tests for General Linear Hypotheses
## 
## Multiple Comparisons of Means: Tukey Contrasts
## 
## 
## Fit: aov(formula = MCBN ~ Group + Alcohol + Marj + Age + Dataset + 
##     CPZ_eqiuv, data = dataset)
## 
## Linear Hypotheses:
##                  Estimate Std. Error t value Pr(>|t|)  
## CHR - HC == 0  -0.0009782  0.0228613  -0.043   0.9659  
## ECP - HC == 0  -0.0193331  0.0243869  -0.793   0.6425  
## CP - HC == 0   -0.0749526  0.0303706  -2.468   0.0839 .
## ECP - CHR == 0 -0.0183550  0.0337260  -0.544   0.7039  
## CP - CHR == 0  -0.0739744  0.0377878  -1.958   0.1528  
## CP - ECP == 0  -0.0556194  0.0345726  -1.609   0.2168  
## ---
## Signif. codes:  0 '***' 0.001 '**' 0.01 '*' 0.05 '.' 0.1 ' ' 1
## (Adjusted p values reported -- fdr method)
```

### Group Differences in Cort GE

#### FPN

```
GroupDiffFPN <- aov(FPN ~ Group + Alcohol+ Marj + Age + Dataset + CPZ_eqiuv, dataset)
summary(GroupDiffFPN)
```

```
##              Df Sum Sq  Mean Sq F value Pr(>F)
## Group         3 0.0251 0.008358   1.175  0.319
## Alcohol       1 0.0106 0.010565   1.486  0.224
## Marj          1 0.0104 0.010354   1.456  0.228
## Age           1 0.0112 0.011224   1.578  0.210
## Dataset       2 0.0196 0.009815   1.380  0.253
## CPZ_eqiuv     1 0.0016 0.001613   0.227  0.634
## Residuals   428 3.0438 0.007112               
## 10 observations deleted due to missingness
```

```
FPN_group_plot <- effect_plot(GroupDiffFPN, pred = Group, interval = TRUE, partial.residuals = TRUE, jitter = .2)
FPN_group_plot + theme(text = element_text(size = 24), panel.border = element_rect(colour = "black", fill=NA, size=1), axis.title.x = element_blank())
```

```
posthoc_FPN <- glht(GroupDiffFPN, linfct = mcp(Group="Tukey"))
summary(posthoc_FPN, adjusted(type='fdr'))
```

```
## 
##   Simultaneous Tests for General Linear Hypotheses
## 
## Multiple Comparisons of Means: Tukey Contrasts
## 
## 
## Fit: aov(formula = FPN ~ Group + Alcohol + Marj + Age + Dataset + 
##     CPZ_eqiuv, data = dataset)
## 
## Linear Hypotheses:
##                 Estimate Std. Error t value Pr(>|t|)
## CHR - HC == 0   0.004362   0.013120   0.332    0.888
## ECP - HC == 0  -0.008782   0.013996  -0.628    0.888
## CP - HC == 0    0.007336   0.017430   0.421    0.888
## ECP - CHR == 0 -0.013145   0.019355  -0.679    0.888
## CP - CHR == 0   0.002974   0.021686   0.137    0.891
## CP - ECP == 0   0.016118   0.019841   0.812    0.888
## (Adjusted p values reported -- fdr method)
```

#### CON

```
GroupDiffCON <- aov(CON ~ Group + Alcohol+ Marj + Age + Dataset + CPZ_eqiuv, dataset)
summary(GroupDiffCON)
```

```
##              Df Sum Sq Mean Sq F value Pr(>F)  
## Group         3  0.256 0.08543   3.813 0.0102 *
## Alcohol       1  0.015 0.01527   0.681 0.4095  
## Marj          1  0.003 0.00306   0.137 0.7119  
## Age           1  0.000 0.00000   0.000 0.9945  
## Dataset       2  0.096 0.04793   2.139 0.1190  
## CPZ_eqiuv     1  0.000 0.00000   0.000 0.9951  
## Residuals   428  9.590 0.02241                 
## ---
## Signif. codes:  0 '***' 0.001 '**' 0.01 '*' 0.05 '.' 0.1 ' ' 1
## 10 observations deleted due to missingness
```

```
CON_group_plot <- effect_plot(GroupDiffCON, pred = Group, interval = TRUE, partial.residuals = TRUE, jitter = .2)
CON_group_plot + theme(text = element_text(size = 24), panel.border = element_rect(colour = "black", fill=NA, size=1), axis.title.x = element_blank())
```

```
posthoc_CON <- glht(GroupDiffCON, linfct = mcp(Group="Tukey"))
summary(posthoc_CON, adjusted(type='fdr'))
```

```
## 
##   Simultaneous Tests for General Linear Hypotheses
## 
## Multiple Comparisons of Means: Tukey Contrasts
## 
## 
## Fit: aov(formula = CON ~ Group + Alcohol + Marj + Age + Dataset + 
##     CPZ_eqiuv, data = dataset)
## 
## Linear Hypotheses:
##                 Estimate Std. Error t value Pr(>|t|)
## CHR - HC == 0  -0.050939   0.023289  -2.187    0.176
## ECP - HC == 0  -0.025262   0.024843  -1.017    0.491
## CP - HC == 0    0.009255   0.030938   0.299    0.765
## ECP - CHR == 0  0.025677   0.034356   0.747    0.546
## CP - CHR == 0   0.060194   0.038494   1.564    0.356
## CP - ECP == 0   0.034517   0.035219   0.980    0.491
## (Adjusted p values reported -- fdr method)
```

#### DMN

```
GroupDiffDMN <- aov(DMN ~ Group + Alcohol+ Marj + Age + Dataset + CPZ_eqiuv, dataset)
summary(GroupDiffDMN)
```

```
##              Df Sum Sq Mean Sq F value  Pr(>F)   
## Group         3 0.1076 0.03586   4.898 0.00234 **
## Alcohol       1 0.0001 0.00008   0.011 0.91671   
## Marj          1 0.0028 0.00279   0.381 0.53748   
## Age           1 0.0065 0.00654   0.894 0.34498   
## Dataset       2 0.0243 0.01216   1.661 0.19124   
## CPZ_eqiuv     1 0.0040 0.00404   0.552 0.45778   
## Residuals   428 3.1331 0.00732                   
## ---
## Signif. codes:  0 '***' 0.001 '**' 0.01 '*' 0.05 '.' 0.1 ' ' 1
## 10 observations deleted due to missingness
```

```
DMN_group_plot <- effect_plot(GroupDiffDMN, pred = Group, interval = TRUE, partial.residuals = TRUE, jitter = .2)
DMN_group_plot + theme(text = element_text(size = 24), panel.border = element_rect(colour = "black", fill=NA, size=1), axis.title.x = element_blank())
```

```
posthoc_DMN <- glht(GroupDiffDMN, linfct = mcp(Group="Tukey"))
summary(posthoc_DMN, adjusted(type='fdr'))
```

```
## 
##   Simultaneous Tests for General Linear Hypotheses
## 
## Multiple Comparisons of Means: Tukey Contrasts
## 
## 
## Fit: aov(formula = DMN ~ Group + Alcohol + Marj + Age + Dataset + 
##     CPZ_eqiuv, data = dataset)
## 
## Linear Hypotheses:
##                 Estimate Std. Error t value Pr(>|t|)
## CHR - HC == 0  -0.002882   0.013311  -0.216    0.869
## ECP - HC == 0  -0.025542   0.014200  -1.799    0.437
## CP - HC == 0   -0.006509   0.017684  -0.368    0.869
## ECP - CHR == 0 -0.022661   0.019637  -1.154    0.690
## CP - CHR == 0  -0.003628   0.022002  -0.165    0.869
## CP - ECP == 0   0.019033   0.020130   0.945    0.690
## (Adjusted p values reported -- fdr method)
```

## EN

```
GroupDiffEN <- aov(EN ~ Group + Alcohol+ Marj + Age + Dataset + CPZ_eqiuv, dataset)
summary(GroupDiffEN)
```

```
##              Df Sum Sq  Mean Sq F value Pr(>F)   
## Group         3 0.0846 0.028203   5.448 0.0011 **
## Alcohol       1 0.0000 0.000018   0.003 0.9533   
## Marj          1 0.0115 0.011514   2.224 0.1366   
## Age           1 0.0119 0.011913   2.301 0.1300   
## Dataset       2 0.0224 0.011218   2.167 0.1158   
## CPZ_eqiuv     1 0.0010 0.001027   0.198 0.6563   
## Residuals   428 2.2157 0.005177                  
## ---
## Signif. codes:  0 '***' 0.001 '**' 0.01 '*' 0.05 '.' 0.1 ' ' 1
## 10 observations deleted due to missingness
```

```
EN_group_plot <- effect_plot(GroupDiffEN, pred = Group, interval = TRUE, partial.residuals = TRUE, jitter = .2)
EN_group_plot + theme(text = element_text(size = 24), panel.border = element_rect(colour = "black", fill=NA, size=1), axis.title.x = element_blank())
```

```
posthoc_EN <- glht(GroupDiffEN, linfct = mcp(Group="Tukey"))
summary(posthoc_EN, adjusted(type='fdr'))
```

```
## 
##   Simultaneous Tests for General Linear Hypotheses
## 
## Multiple Comparisons of Means: Tukey Contrasts
## 
## 
## Fit: aov(formula = EN ~ Group + Alcohol + Marj + Age + Dataset + CPZ_eqiuv, 
##     data = dataset)
## 
## Linear Hypotheses:
##                  Estimate Std. Error t value Pr(>|t|)
## CHR - HC == 0  -0.0238427  0.0111941  -2.130    0.202
## ECP - HC == 0  -0.0115745  0.0119411  -0.969    0.609
## CP - HC == 0   -0.0003426  0.0148710  -0.023    0.982
## ECP - CHR == 0  0.0122682  0.0165140   0.743    0.609
## CP - CHR == 0   0.0235001  0.0185029   1.270    0.609
## CP - ECP == 0   0.0112319  0.0169286   0.663    0.609
## (Adjusted p values reported -- fdr method)
```

## MN

```
GroupDiffMN <- aov(MN ~ Group + Alcohol+ Marj + Age + Dataset + CPZ_eqiuv, dataset)
summary(GroupDiffMN)
```

```
##              Df Sum Sq Mean Sq F value   Pr(>F)    
## Group         3 0.0269 0.00898   1.645    0.178    
## Alcohol       1 0.0001 0.00012   0.022    0.882    
## Marj          1 0.0037 0.00371   0.680    0.410    
## Age           1 0.0000 0.00001   0.002    0.966    
## Dataset       2 0.1950 0.09749  17.864 3.54e-08 ***
## CPZ_eqiuv     1 0.0033 0.00329   0.603    0.438    
## Residuals   427 2.3301 0.00546                     
## ---
## Signif. codes:  0 '***' 0.001 '**' 0.01 '*' 0.05 '.' 0.1 ' ' 1
## 11 observations deleted due to missingness
```

```
MN_group_plot <- effect_plot(GroupDiffMN, pred = Group, interval = TRUE, partial.residuals = TRUE, jitter = .2)
MN_group_plot + theme(text = element_text(size = 24), panel.border = element_rect(colour = "black", fill=NA, size=1), axis.title.x = element_blank())
```

```
posthoc_MN <- glht(GroupDiffMN, linfct = mcp(Group="Tukey"))
summary(posthoc_MN, adjusted(type='fdr'))
```

```
## 
##   Simultaneous Tests for General Linear Hypotheses
## 
## Multiple Comparisons of Means: Tukey Contrasts
## 
## 
## Fit: aov(formula = MN ~ Group + Alcohol + Marj + Age + Dataset + CPZ_eqiuv, 
##     data = dataset)
## 
## Linear Hypotheses:
##                 Estimate Std. Error t value Pr(>|t|)
## CHR - HC == 0  -0.005410   0.011526  -0.469    0.725
## ECP - HC == 0   0.004321   0.012260   0.352    0.725
## CP - HC == 0   -0.016202   0.015268  -1.061    0.725
## ECP - CHR == 0  0.009731   0.016978   0.573    0.725
## CP - CHR == 0  -0.010791   0.019018  -0.567    0.725
## CP - ECP == 0  -0.020523   0.017380  -1.181    0.725
## (Adjusted p values reported -- fdr method)
```

## AN

```
GroupDiffAN <- aov(AN ~ Group + Alcohol+ Marj + Age + Dataset + CPZ_eqiuv, dataset)
summary(GroupDiffAN)
```

```
##              Df Sum Sq Mean Sq F value   Pr(>F)    
## Group         3 0.0837 0.02791   4.427 0.004440 ** 
## Alcohol       1 0.0190 0.01902   3.016 0.083166 .  
## Marj          1 0.0028 0.00280   0.445 0.505228    
## Age           1 0.0259 0.02594   4.115 0.043132 *  
## Dataset       2 0.1092 0.05459   8.658 0.000206 ***
## CPZ_eqiuv     1 0.0034 0.00336   0.534 0.465492    
## Residuals   428 2.6987 0.00631                     
## ---
## Signif. codes:  0 '***' 0.001 '**' 0.01 '*' 0.05 '.' 0.1 ' ' 1
## 10 observations deleted due to missingness
```

```
AN_group_plot <- effect_plot(GroupDiffAN, pred = Group, interval = TRUE, partial.residuals = TRUE, jitter = .2)
AN_group_plot + theme(text = element_text(size = 24), panel.border = element_rect(colour = "black", fill=NA, size=1), axis.title.x = element_blank())
```

```
posthoc_AN <- glht(GroupDiffAN, linfct = mcp(Group="Tukey"))
summary(posthoc_AN, adjusted(type='fdr'))
```

```
## 
##   Simultaneous Tests for General Linear Hypotheses
## 
## Multiple Comparisons of Means: Tukey Contrasts
## 
## 
## Fit: aov(formula = AN ~ Group + Alcohol + Marj + Age + Dataset + CPZ_eqiuv, 
##     data = dataset)
## 
## Linear Hypotheses:
##                 Estimate Std. Error t value Pr(>|t|)
## CHR - HC == 0  -0.020031   0.012354  -1.621    0.211
## ECP - HC == 0  -0.025840   0.013178  -1.961    0.152
## CP - HC == 0   -0.038306   0.016412  -2.334    0.120
## ECP - CHR == 0 -0.005809   0.018225  -0.319    0.750
## CP - CHR == 0  -0.018275   0.020420  -0.895    0.557
## CP - ECP == 0  -0.012466   0.018683  -0.667    0.606
## (Adjusted p values reported -- fdr method)
```

## VN

```
GroupDiffVN <- aov(VN ~ Group + Alcohol+ Marj + Age + Dataset + CPZ_eqiuv, dataset)
summary(GroupDiffVN)
```

```
##              Df Sum Sq  Mean Sq F value Pr(>F)  
## Group         3 0.0339 0.011294   3.336 0.0194 *
## Alcohol       1 0.0100 0.010023   2.960 0.0861 .
## Marj          1 0.0029 0.002917   0.862 0.3538  
## Age           1 0.0095 0.009529   2.815 0.0941 .
## Dataset       2 0.0027 0.001357   0.401 0.6700  
## CPZ_eqiuv     1 0.0058 0.005805   1.715 0.1911  
## Residuals   428 1.4491 0.003386                 
## ---
## Signif. codes:  0 '***' 0.001 '**' 0.01 '*' 0.05 '.' 0.1 ' ' 1
## 10 observations deleted due to missingness
```

```
VN_group_plot <- effect_plot(GroupDiffVN, pred = Group, interval = TRUE, partial.residuals = TRUE, jitter = .2)
VN_group_plot + theme(text = element_text(size = 24), panel.border = element_rect(colour = "black", fill=NA, size=1), axis.title.x = element_blank())
```

```
posthoc_VN <- glht(GroupDiffVN, linfct = mcp(Group="Tukey"))
summary(posthoc_VN, adjusted(type='fdr'))
```

```
## 
##   Simultaneous Tests for General Linear Hypotheses
## 
## Multiple Comparisons of Means: Tukey Contrasts
## 
## 
## Fit: aov(formula = VN ~ Group + Alcohol + Marj + Age + Dataset + CPZ_eqiuv, 
##     data = dataset)
## 
## Linear Hypotheses:
##                 Estimate Std. Error t value Pr(>|t|)
## CHR - HC == 0  -0.002999   0.009053  -0.331    0.864
## ECP - HC == 0  -0.005282   0.009657  -0.547    0.864
## CP - HC == 0   -0.018934   0.012026  -1.574    0.638
## ECP - CHR == 0 -0.002283   0.013355  -0.171    0.864
## CP - CHR == 0  -0.015935   0.014963  -1.065    0.638
## CP - ECP == 0  -0.013653   0.013690  -0.997    0.638
## (Adjusted p values reported -- fdr method)
```

### CB-BG GE predicts Cort GE

#### FPN

```
FPN_4groups <- lm(FPN ~ CCBN*MCBN*Group + Alcohol+ Marj + Age + Dataset + CPZ_eqiuv, data = dataset)
summary(FPN_4groups)
```

```
## 
## Call:
## lm(formula = FPN ~ CCBN * MCBN * Group + Alcohol + Marj + Age + 
##     Dataset + CPZ_eqiuv, data = dataset)
## 
## Residuals:
##       Min        1Q    Median        3Q       Max 
## -0.300131 -0.047405  0.005178  0.052240  0.268794 
## 
## Coefficients:
##                      Estimate Std. Error t value Pr(>|t|)  
## (Intercept)         4.320e-01  1.686e-01   2.562   0.0108 *
## CCBN                3.379e-01  2.350e-01   1.438   0.1512  
## MCBN                3.323e-01  2.608e-01   1.274   0.2033  
## GroupCHR           -6.662e-02  2.904e-01  -0.229   0.8187  
## GroupECP           -1.341e-01  3.687e-01  -0.364   0.7163  
## GroupCP            -3.046e-01  5.176e-01  -0.588   0.5566  
## Alcohol            -9.752e-03  7.536e-03  -1.294   0.1964  
## Marj                1.231e-02  7.088e-03   1.737   0.0832 .
## Age                -7.555e-04  5.757e-04  -1.312   0.1901  
## DatasetCOBRE       -1.213e-03  1.969e-02  -0.062   0.9509  
## DatasetHCP         -1.228e-02  2.032e-02  -0.604   0.5461  
## CPZ_eqiuv          -3.523e-06  7.464e-06  -0.472   0.6372  
## CCBN:MCBN          -3.727e-01  3.577e-01  -1.042   0.2981  
## CCBN:GroupCHR       1.559e-01  4.255e-01   0.366   0.7143  
## CCBN:GroupECP       1.495e-01  5.216e-01   0.287   0.7746  
## CCBN:GroupCP        6.009e-01  7.325e-01   0.820   0.4125  
## MCBN:GroupCHR      -1.721e-02  4.493e-01  -0.038   0.9695  
## MCBN:GroupECP       4.814e-01  5.776e-01   0.833   0.4051  
## MCBN:GroupCP        5.229e-01  7.608e-01   0.687   0.4923  
## CCBN:MCBN:GroupCHR -4.283e-02  6.394e-01  -0.067   0.9466  
## CCBN:MCBN:GroupECP -6.279e-01  8.043e-01  -0.781   0.4354  
## CCBN:MCBN:GroupCP  -9.606e-01  1.067e+00  -0.901   0.3683  
## ---
## Signif. codes:  0 '***' 0.001 '**' 0.01 '*' 0.05 '.' 0.1 ' ' 1
## 
## Residual standard error: 0.08304 on 416 degrees of freedom
##   (10 observations deleted due to missingness)
## Multiple R-squared:  0.0813, Adjusted R-squared:  0.03493 
## F-statistic: 1.753 on 21 and 416 DF,  p-value: 0.02146
```

```
FPN_CCBN_plot <- effect_plot(FPN_4groups, pred = CCBN, interval = TRUE, partial.residuals = TRUE)
FPN_CCBN_plot + theme(text = element_text(size = 24), panel.border = element_rect(colour = "black", fill=NA, size=1))
```

```
FPN_MCBN_plot <- effect_plot(FPN_4groups, pred = MCBN, interval = TRUE, partial.residuals = TRUE)
FPN_MCBN_plot + theme(text = element_text(size = 24), panel.border = element_rect(colour = "black", fill=NA, size=1))
```

```
trellis.par.set(par.axis.text=list(cex=1.25))
trellis.par.set(par.strip.text=list(cex=2))
trellis.par.set(par.xlab.text=list(fontsize=0))
trellis.par.set(par.ylab.text=list(fontsize=0))

xyplot(FPN ~ CCBN | Group, data=dataset, fit = FPN_4groups, par.settings = list(strip.background=list(col="lightgrey"), par.strip.text = list(fontsize=20)),
  panel = function(x, y, ...) {
       panel.xyplot(x, y, ..., col = "black")
       panel.lmline(x, y, col = "black")
  })
```

```
xyplot(FPN ~ MCBN | Group, data=dataset, par.settings = list(strip.background=list(col="lightgrey")), 
  panel = function(x, y, ...) {
       panel.xyplot(x, y, ..., col = "black")
       panel.lmline(x, y, col = "black")
  })
```

#### CON

```
CON_4groups <- lm(CON ~ CCBN*MCBN*Group + Alcohol+ Marj + Age + Dataset + CPZ_eqiuv, data = dataset)
summary(CON_4groups)
```

```
## 
## Call:
## lm(formula = CON ~ CCBN * MCBN * Group + Alcohol + Marj + Age + 
##     Dataset + CPZ_eqiuv, data = dataset)
## 
## Residuals:
##      Min       1Q   Median       3Q      Max 
## -0.45138 -0.08449  0.02508  0.09998  0.30597 
## 
## Coefficients:
##                      Estimate Std. Error t value Pr(>|t|)  
## (Intercept)         1.995e-01  3.016e-01   0.662   0.5086  
## CCBN                9.312e-01  4.204e-01   2.215   0.0273 *
## MCBN                7.071e-01  4.666e-01   1.515   0.1304  
## GroupCHR            3.645e-01  5.195e-01   0.702   0.4833  
## GroupECP           -1.901e-02  6.594e-01  -0.029   0.9770  
## GroupCP             8.264e-02  9.259e-01   0.089   0.9289  
## Alcohol            -1.340e-02  1.348e-02  -0.994   0.3207  
## Marj                8.232e-04  1.268e-02   0.065   0.9483  
## Age                -3.891e-04  1.030e-03  -0.378   0.7058  
## DatasetCOBRE        2.025e-02  3.521e-02   0.575   0.5656  
## DatasetHCP         -1.785e-02  3.635e-02  -0.491   0.6236  
## CPZ_eqiuv           2.141e-06  1.335e-05   0.160   0.8727  
## CCBN:MCBN          -1.188e+00  6.399e-01  -1.856   0.0641 .
## CCBN:GroupCHR      -7.454e-01  7.612e-01  -0.979   0.3281  
## CCBN:GroupECP      -2.665e-01  9.330e-01  -0.286   0.7753  
## CCBN:GroupCP       -4.151e-02  1.310e+00  -0.032   0.9747  
## MCBN:GroupCHR      -7.482e-01  8.037e-01  -0.931   0.3524  
## MCBN:GroupECP      -5.053e-02  1.033e+00  -0.049   0.9610  
## MCBN:GroupCP       -2.224e-01  1.361e+00  -0.163   0.8702  
## CCBN:MCBN:GroupCHR  1.290e+00  1.144e+00   1.128   0.2602  
## CCBN:MCBN:GroupECP  4.505e-01  1.439e+00   0.313   0.7544  
## CCBN:MCBN:GroupCP   2.139e-01  1.908e+00   0.112   0.9108  
## ---
## Signif. codes:  0 '***' 0.001 '**' 0.01 '*' 0.05 '.' 0.1 ' ' 1
## 
## Residual standard error: 0.1485 on 416 degrees of freedom
##   (10 observations deleted due to missingness)
## Multiple R-squared:  0.07854,    Adjusted R-squared:  0.03203 
## F-statistic: 1.689 on 21 and 416 DF,  p-value: 0.02972
```

```
CON_CCBN_plot <- effect_plot(CON_4groups, pred = CCBN, interval = TRUE, partial.residuals = TRUE)
CON_CCBN_plot + theme(text = element_text(size = 24), panel.border = element_rect(colour = "black", fill=NA, size=1))
```

```
CON_MCBN_plot <- effect_plot(CON_4groups, pred = MCBN, interval = TRUE, partial.residuals = TRUE)
CON_MCBN_plot + theme(text = element_text(size = 24), panel.border = element_rect(colour = "black", fill=NA, size=1))
```

```
xyplot(FPN ~ CCBN | Group, data=dataset, par.settings = list(strip.background=list(col="lightgrey")), 
  panel = function(x, y, ...) {
       panel.xyplot(x, y, ..., col = "black")
       panel.lmline(x, y, col = "black")
  })
```

```
xyplot(FPN ~ MCBN | Group, data=dataset, par.settings = list(strip.background=list(col="lightgrey")), 
  panel = function(x, y, ...) {
       panel.xyplot(x, y, ..., col = "black")
       panel.lmline(x, y, col = "black")
  })
```

#### DMN

```
DMN_4groups <- lm(DMN ~ CCBN*MCBN*Group + Alcohol+ Marj + Age + Dataset + CPZ_eqiuv, data = dataset)
summary(DMN_4groups)
```

```
## 
## Call:
## lm(formula = DMN ~ CCBN * MCBN * Group + Alcohol + Marj + Age + 
##     Dataset + CPZ_eqiuv, data = dataset)
## 
## Residuals:
##       Min        1Q    Median        3Q       Max 
## -0.289668 -0.050216  0.008165  0.052942  0.272563 
## 
## Coefficients:
##                      Estimate Std. Error t value Pr(>|t|)    
## (Intercept)         6.399e-01  1.732e-01   3.695 0.000249 ***
## CCBN                1.569e-01  2.414e-01   0.650 0.515933    
## MCBN                1.829e-01  2.679e-01   0.683 0.495112    
## GroupCHR            1.290e-01  2.983e-01   0.432 0.665679    
## GroupECP           -8.584e-01  3.786e-01  -2.267 0.023883 *  
## GroupCP             5.723e-01  5.316e-01   1.077 0.282283    
## Alcohol             1.157e-03  7.740e-03   0.149 0.881267    
## Marj                9.445e-03  7.279e-03   1.298 0.195170    
## Age                -5.021e-04  5.912e-04  -0.849 0.396212    
## DatasetCOBRE       -1.211e-02  2.022e-02  -0.599 0.549492    
## DatasetHCP         -2.898e-02  2.087e-02  -1.388 0.165764    
## CPZ_eqiuv          -5.693e-06  7.666e-06  -0.743 0.458156    
## CCBN:MCBN          -2.450e-01  3.674e-01  -0.667 0.505302    
## CCBN:GroupCHR      -1.841e-01  4.370e-01  -0.421 0.673734    
## CCBN:GroupECP       1.124e+00  5.357e-01   2.099 0.036408 *  
## CCBN:GroupCP       -7.960e-01  7.523e-01  -1.058 0.290669    
## MCBN:GroupCHR      -1.332e-01  4.614e-01  -0.289 0.773012    
## MCBN:GroupECP       1.124e+00  5.932e-01   1.895 0.058801 .  
## MCBN:GroupCP       -1.052e+00  7.814e-01  -1.347 0.178866    
## CCBN:MCBN:GroupCHR  1.820e-01  6.566e-01   0.277 0.781732    
## CCBN:MCBN:GroupECP -1.503e+00  8.261e-01  -1.819 0.069637 .  
## CCBN:MCBN:GroupCP   1.451e+00  1.095e+00   1.325 0.185900    
## ---
## Signif. codes:  0 '***' 0.001 '**' 0.01 '*' 0.05 '.' 0.1 ' ' 1
## 
## Residual standard error: 0.08528 on 416 degrees of freedom
##   (10 observations deleted due to missingness)
## Multiple R-squared:  0.07715,    Adjusted R-squared:  0.03056 
## F-statistic: 1.656 on 21 and 416 DF,  p-value: 0.03488
```

```
DMN_CCBN_plot <- effect_plot(DMN_4groups, pred = CCBN, interval = TRUE, partial.residuals = TRUE)
DMN_CCBN_plot + theme(text = element_text(size = 24), panel.border = element_rect(colour = "black", fill=NA, size=1))
```

```
DMN_MCBN_plot <- effect_plot(DMN_4groups, pred = MCBN, interval = TRUE, partial.residuals = TRUE)
DMN_MCBN_plot + theme(text = element_text(size = 24), panel.border = element_rect(colour = "black", fill=NA, size=1))
```

```
xyplot(DMN ~ CCBN | Group, data=dataset, par.settings = list(strip.background=list(col="lightgrey")), 
  panel = function(x, y, ...) {
       panel.xyplot(x, y, ..., col = "black")
       panel.lmline(x, y, col = "black")
  })
```

```
xyplot(DMN ~ MCBN | Group, data=dataset, par.settings = list(strip.background=list(col="lightgrey")), 
  panel = function(x, y, ...) {
       panel.xyplot(x, y, ..., col = "black")
       panel.lmline(x, y, col = "black")
  })
```

## EN

```
EN_4groups <- lm(EN ~ CCBN*MCBN*Group + Alcohol+ Marj + Age + Dataset + CPZ_eqiuv, data = dataset)
summary(EN_4groups)
```

```
## 
## Call:
## lm(formula = EN ~ CCBN * MCBN * Group + Alcohol + Marj + Age + 
##     Dataset + CPZ_eqiuv, data = dataset)
## 
## Residuals:
##      Min       1Q   Median       3Q      Max 
## -0.33919 -0.03767  0.00706  0.04532  0.18161 
## 
## Coefficients:
##                      Estimate Std. Error t value Pr(>|t|)  
## (Intercept)         3.671e-01  1.442e-01   2.547   0.0112 *
## CCBN                3.347e-01  2.009e-01   1.666   0.0965 .
## MCBN                3.225e-01  2.230e-01   1.446   0.1488  
## GroupCHR            2.233e-01  2.483e-01   0.899   0.3690  
## GroupECP           -7.594e-03  3.152e-01  -0.024   0.9808  
## GroupCP             2.583e-01  4.426e-01   0.584   0.5598  
## Alcohol            -7.608e-04  6.443e-03  -0.118   0.9061  
## Marj               -9.596e-03  6.060e-03  -1.584   0.1140  
## Age                 4.443e-05  4.922e-04   0.090   0.9281  
## DatasetCOBRE        2.714e-02  1.683e-02   1.612   0.1077  
## DatasetHCP          2.161e-02  1.738e-02   1.244   0.2142  
## CPZ_eqiuv          -3.401e-06  6.382e-06  -0.533   0.5944  
## CCBN:MCBN          -3.703e-01  3.059e-01  -1.211   0.2267  
## CCBN:GroupCHR      -3.629e-01  3.638e-01  -0.998   0.3191  
## CCBN:GroupECP      -1.331e-02  4.459e-01  -0.030   0.9762  
## CCBN:GroupCP       -2.770e-01  6.263e-01  -0.442   0.6585  
## MCBN:GroupCHR      -4.460e-01  3.841e-01  -1.161   0.2462  
## MCBN:GroupECP       2.060e-01  4.938e-01   0.417   0.6767  
## MCBN:GroupCP       -1.344e-01  6.505e-01  -0.207   0.8364  
## CCBN:MCBN:GroupCHR  6.511e-01  5.466e-01   1.191   0.2343  
## CCBN:MCBN:GroupECP -2.737e-01  6.877e-01  -0.398   0.6908  
## CCBN:MCBN:GroupCP   7.156e-02  9.119e-01   0.078   0.9375  
## ---
## Signif. codes:  0 '***' 0.001 '**' 0.01 '*' 0.05 '.' 0.1 ' ' 1
## 
## Residual standard error: 0.07099 on 416 degrees of freedom
##   (10 observations deleted due to missingness)
## Multiple R-squared:  0.1067, Adjusted R-squared:  0.06165 
## F-statistic: 2.367 on 21 and 416 DF,  p-value: 0.0006845
```

```
EN_CCBN_plot <- effect_plot(EN_4groups, pred = CCBN, interval = TRUE, partial.residuals = TRUE)
EN_CCBN_plot + theme(text = element_text(size = 24), panel.border = element_rect(colour = "black", fill=NA, size=1))
```

```
EN_MCBN_plot <- effect_plot(EN_4groups, pred = MCBN, interval = TRUE, partial.residuals = TRUE)
EN_MCBN_plot + theme(text = element_text(size = 24), panel.border = element_rect(colour = "black", fill=NA, size=1))
```

```
xyplot(EN ~ CCBN | Group, data=dataset, par.settings = list(strip.background=list(col="lightgrey")), 
  panel = function(x, y, ...) {
       panel.xyplot(x, y, ..., col = "black")
       panel.lmline(x, y, col = "black")
  })
```

```
xyplot(EN ~ MCBN | Group, data=dataset, par.settings = list(strip.background=list(col="lightgrey")), 
  panel = function(x, y, ...) {
       panel.xyplot(x, y, ..., col = "black")
       panel.lmline(x, y, col = "black")
  })
```

## MN

```
MN_4groups <- lm(MN ~ CCBN*MCBN*Group + Alcohol+ Marj + Age + Dataset + CPZ_eqiuv, data = dataset)
summary(MN_4groups)
```

```
## 
## Call:
## lm(formula = MN ~ CCBN * MCBN * Group + Alcohol + Marj + Age + 
##     Dataset + CPZ_eqiuv, data = dataset)
## 
## Residuals:
##       Min        1Q    Median        3Q       Max 
## -0.296287 -0.044143 -0.000827  0.046852  0.182739 
## 
## Coefficients:
##                      Estimate Std. Error t value Pr(>|t|)    
## (Intercept)         6.855e-01  1.479e-01   4.635 4.79e-06 ***
## CCBN                1.172e-01  2.061e-01   0.568   0.5700    
## MCBN                9.613e-02  2.288e-01   0.420   0.6746    
## GroupCHR            5.484e-02  2.547e-01   0.215   0.8296    
## GroupECP           -7.252e-01  3.234e-01  -2.243   0.0254 *  
## GroupCP            -5.391e-01  4.540e-01  -1.187   0.2357    
## Alcohol             3.649e-03  6.614e-03   0.552   0.5815    
## Marj                4.119e-03  6.217e-03   0.663   0.5080    
## Age                -9.368e-04  5.050e-04  -1.855   0.0643 .  
## DatasetCOBRE        1.887e-02  1.727e-02   1.093   0.2750    
## DatasetHCP         -4.774e-02  1.783e-02  -2.678   0.0077 ** 
## CPZ_eqiuv           1.638e-06  6.547e-06   0.250   0.8025    
## CCBN:MCBN          -6.828e-02  3.138e-01  -0.218   0.8279    
## CCBN:GroupCHR      -9.792e-02  3.732e-01  -0.262   0.7932    
## CCBN:GroupECP       1.064e+00  4.575e-01   2.325   0.0206 *  
## CCBN:GroupCP        6.518e-01  6.425e-01   1.014   0.3109    
## MCBN:GroupCHR      -1.016e-01  3.941e-01  -0.258   0.7966    
## MCBN:GroupECP       1.257e+00  5.066e-01   2.482   0.0135 *  
## MCBN:GroupCP        7.973e-01  6.673e-01   1.195   0.2328    
## CCBN:MCBN:GroupCHR  1.655e-01  5.609e-01   0.295   0.7681    
## CCBN:MCBN:GroupECP -1.809e+00  7.055e-01  -2.565   0.0107 *  
## CCBN:MCBN:GroupCP  -9.775e-01  9.355e-01  -1.045   0.2967    
## ---
## Signif. codes:  0 '***' 0.001 '**' 0.01 '*' 0.05 '.' 0.1 ' ' 1
## 
## Residual standard error: 0.07283 on 415 degrees of freedom
##   (11 observations deleted due to missingness)
## Multiple R-squared:  0.1398, Adjusted R-squared:  0.09623 
## F-statistic: 3.211 on 21 and 415 DF,  p-value: 3.246e-06
```

```
MN_CCBN_plot <- effect_plot(MN_4groups, pred = CCBN, interval = TRUE, partial.residuals = TRUE)
MN_CCBN_plot + theme(text = element_text(size = 24), panel.border = element_rect(colour = "black", fill=NA, size=1))
```

```
MN_MCBN_plot <- effect_plot(MN_4groups, pred = MCBN, interval = TRUE, partial.residuals = TRUE)
MN_MCBN_plot + theme(text = element_text(size = 24), panel.border = element_rect(colour = "black", fill=NA, size=1))
```

```
xyplot(MN ~ CCBN | Group, data=dataset, par.settings = list(strip.background=list(col="lightgrey")), 
  panel = function(x, y, ...) {
       panel.xyplot(x, y, ..., col = "black")
       panel.lmline(x, y, col = "black")
  })
```

```
xyplot(MN ~ MCBN | Group, data=dataset, par.settings = list(strip.background=list(col="lightgrey")), 
  panel = function(x, y, ...) {
       panel.xyplot(x, y, ..., col = "black")
       panel.lmline(x, y, col = "black")
  })
```

## VN

```
VN_4groups <- lm(VN ~ CCBN*MCBN*Group + Alcohol+ Marj + Age + Dataset + CPZ_eqiuv, data = dataset)
summary(VN_4groups)
```

```
## 
## Call:
## lm(formula = VN ~ CCBN * MCBN * Group + Alcohol + Marj + Age + 
##     Dataset + CPZ_eqiuv, data = dataset)
## 
## Residuals:
##       Min        1Q    Median        3Q       Max 
## -0.149975 -0.036045 -0.002834  0.035920  0.138783 
## 
## Coefficients:
##                      Estimate Std. Error t value Pr(>|t|)    
## (Intercept)         9.512e-01  1.174e-01   8.105 5.95e-15 ***
## CCBN               -1.931e-01  1.636e-01  -1.181  0.23840    
## MCBN               -1.792e-01  1.815e-01  -0.987  0.32407    
## GroupCHR           -2.012e-01  2.021e-01  -0.995  0.32018    
## GroupECP           -8.079e-01  2.566e-01  -3.149  0.00176 ** 
## GroupCP            -2.434e-01  3.603e-01  -0.676  0.49967    
## Alcohol            -1.445e-03  5.245e-03  -0.275  0.78308    
## Marj               -4.507e-03  4.933e-03  -0.914  0.36140    
## Age                -3.383e-04  4.007e-04  -0.844  0.39902    
## DatasetCOBRE       -9.058e-03  1.370e-02  -0.661  0.50891    
## DatasetHCP          2.482e-03  1.415e-02   0.175  0.86080    
## CPZ_eqiuv           5.277e-06  5.195e-06   1.016  0.31034    
## CCBN:MCBN           2.715e-01  2.490e-01   1.091  0.27610    
## CCBN:GroupCHR       3.835e-01  2.962e-01   1.295  0.19613    
## CCBN:GroupECP       1.105e+00  3.630e-01   3.043  0.00249 ** 
## CCBN:GroupCP        3.538e-01  5.099e-01   0.694  0.48816    
## MCBN:GroupCHR       3.084e-01  3.127e-01   0.986  0.32464    
## MCBN:GroupECP       1.286e+00  4.020e-01   3.200  0.00148 ** 
## MCBN:GroupCP        3.427e-01  5.295e-01   0.647  0.51789    
## CCBN:MCBN:GroupCHR -5.792e-01  4.450e-01  -1.302  0.19380    
## CCBN:MCBN:GroupECP -1.757e+00  5.598e-01  -3.139  0.00182 ** 
## CCBN:MCBN:GroupCP  -5.299e-01  7.423e-01  -0.714  0.47575    
## ---
## Signif. codes:  0 '***' 0.001 '**' 0.01 '*' 0.05 '.' 0.1 ' ' 1
## 
## Residual standard error: 0.05779 on 416 degrees of freedom
##   (10 observations deleted due to missingness)
## Multiple R-squared:  0.08217,    Adjusted R-squared:  0.03583 
## F-statistic: 1.773 on 21 and 416 DF,  p-value: 0.01933
```

```
VN_CCBN_plot <- effect_plot(VN_4groups, pred = CCBN, interval = TRUE, partial.residuals = TRUE)
VN_CCBN_plot + theme(text = element_text(size = 24), panel.border = element_rect(colour = "black", fill=NA, size=1))
```

```
VN_MCBN_plot <- effect_plot(VN_4groups, pred = MCBN, interval = TRUE, partial.residuals = TRUE)
VN_MCBN_plot + theme(text = element_text(size = 24), panel.border = element_rect(colour = "black", fill=NA, size=1))
```

```
xyplot(VN ~ CCBN | Group, data=dataset, par.settings = list(strip.background=list(col="lightgrey")), 
  panel = function(x, y, ...) {
       panel.xyplot(x, y, ..., col = "black")
       panel.lmline(x, y, col = "black")
  })
```

```
xyplot(VN ~ MCBN | Group, data=dataset, par.settings = list(strip.background=list(col="lightgrey")), 
  panel = function(x, y, ...) {
       panel.xyplot(x, y, ..., col = "black")
       panel.lmline(x, y, col = "black")
  })
```

## AN

```
AN_4groups <- lm(AN ~ CCBN*MCBN*Group + Alcohol+ Marj + Age + Dataset + CPZ_eqiuv, data = dataset)
summary(AN_4groups)
```

```
## 
## Call:
## lm(formula = AN ~ CCBN * MCBN * Group + Alcohol + Marj + Age + 
##     Dataset + CPZ_eqiuv, data = dataset)
## 
## Residuals:
##       Min        1Q    Median        3Q       Max 
## -0.291869 -0.046590  0.003852  0.049693  0.190807 
## 
## Coefficients:
##                      Estimate Std. Error t value Pr(>|t|)   
## (Intercept)         3.772e-01  1.557e-01   2.422  0.01584 * 
## CCBN                5.344e-01  2.170e-01   2.463  0.01420 * 
## MCBN                5.020e-01  2.409e-01   2.084  0.03776 * 
## GroupCHR            3.413e-01  2.682e-01   1.273  0.20390   
## GroupECP           -1.630e-01  3.404e-01  -0.479  0.63235   
## GroupCP            -1.624e-01  4.780e-01  -0.340  0.73428   
## Alcohol             1.415e-02  6.959e-03   2.034  0.04258 * 
## Marj                4.048e-03  6.545e-03   0.619  0.53654   
## Age                -1.473e-03  5.316e-04  -2.771  0.00585 **
## DatasetCOBRE        8.476e-03  1.818e-02   0.466  0.64124   
## DatasetHCP         -2.610e-02  1.877e-02  -1.391  0.16499   
## CPZ_eqiuv           2.743e-06  6.893e-06   0.398  0.69085   
## CCBN:MCBN          -6.043e-01  3.303e-01  -1.829  0.06806 . 
## CCBN:GroupCHR      -5.442e-01  3.929e-01  -1.385  0.16680   
## CCBN:GroupECP       1.470e-01  4.816e-01   0.305  0.76033   
## CCBN:GroupCP        1.500e-01  6.764e-01   0.222  0.82457   
## MCBN:GroupCHR      -6.199e-01  4.149e-01  -1.494  0.13586   
## MCBN:GroupECP       3.552e-01  5.333e-01   0.666  0.50583   
## MCBN:GroupCP        1.769e-01  7.025e-01   0.252  0.80130   
## CCBN:MCBN:GroupCHR  9.260e-01  5.904e-01   1.569  0.11752   
## CCBN:MCBN:GroupECP -4.253e-01  7.427e-01  -0.573  0.56717   
## CCBN:MCBN:GroupCP  -1.911e-01  9.849e-01  -0.194  0.84628   
## ---
## Signif. codes:  0 '***' 0.001 '**' 0.01 '*' 0.05 '.' 0.1 ' ' 1
## 
## Residual standard error: 0.07668 on 416 degrees of freedom
##   (10 observations deleted due to missingness)
## Multiple R-squared:  0.1689, Adjusted R-squared:  0.1269 
## F-statistic: 4.025 on 21 and 416 DF,  p-value: 1.347e-08
```

```
AN_CCBN_plot <- effect_plot(AN_4groups, pred = CCBN, interval = TRUE, partial.residuals = TRUE)
AN_CCBN_plot + theme(text = element_text(size = 24), panel.border = element_rect(colour = "black", fill=NA, size=1))
```

```
AN_MCBN_plot <- effect_plot(AN_4groups, pred = MCBN, interval = TRUE, partial.residuals = TRUE)
AN_MCBN_plot + theme(text = element_text(size = 24), panel.border = element_rect(colour = "black", fill=NA, size=1))
```

```
xyplot(AN ~ CCBN | Group, data=dataset, par.settings = list(strip.background=list(col="lightgrey")), 
  panel = function(x, y, ...) {
       panel.xyplot(x, y, ..., col = "black")
       panel.lmline(x, y, col = "black")
  })
```

```
xyplot(AN ~ MCBN | Group, data=dataset, par.settings = list(strip.background=list(col="lightgrey")), 
  panel = function(x, y, ...) {
       panel.xyplot(x, y, ..., col = "black")
       panel.lmline(x, y, col = "black")
  })
```

#### Combined table

```
export_summs(FPN_4groups, CON_4groups,DMN_4groups,EN_4groups,MN_4groups, VN_4groups,AN_4groups, to.file = "docx", file.name = "models.docx", model.names = c("FPN", "CON", "DMN", "EN", "MN", "VN", "AN"))
```

|  | FPN | CON | DMN | EN | MN | VN | AN |
| --- | --- | --- | --- | --- | --- | --- | --- |
| (Intercept) | 0.43 \* | 0.20 | 0.64 \*\*\* | 0.37 \* | 0.69 \*\*\* | 0.95 \*\*\* | 0.38 \* |
|  | (0.17) | (0.30) | (0.17) | (0.14) | (0.15) | (0.12) | (0.16) |
| CCBN | 0.34 | 0.93 \* | 0.16 | 0.33 | 0.12 | -0.19 | 0.53 \* |
|  | (0.24) | (0.42) | (0.24) | (0.20) | (0.21) | (0.16) | (0.22) |
| MCBN | 0.33 | 0.71 | 0.18 | 0.32 | 0.10 | -0.18 | 0.50 \* |
|  | (0.26) | (0.47) | (0.27) | (0.22) | (0.23) | (0.18) | (0.24) |
| GroupCHR | -0.07 | 0.36 | 0.13 | 0.22 | 0.05 | -0.20 | 0.34 |
|  | (0.29) | (0.52) | (0.30) | (0.25) | (0.25) | (0.20) | (0.27) |
| GroupECP | -0.13 | -0.02 | -0.86 \* | -0.01 | -0.73 \* | -0.81 \*\* | -0.16 |
|  | (0.37) | (0.66) | (0.38) | (0.32) | (0.32) | (0.26) | (0.34) |
| GroupCP | -0.30 | 0.08 | 0.57 | 0.26 | -0.54 | -0.24 | -0.16 |
|  | (0.52) | (0.93) | (0.53) | (0.44) | (0.45) | (0.36) | (0.48) |
| Alcohol | -0.01 | -0.01 | 0.00 | -0.00 | 0.00 | -0.00 | 0.01 \* |
|  | (0.01) | (0.01) | (0.01) | (0.01) | (0.01) | (0.01) | (0.01) |
| Marj | 0.01 | 0.00 | 0.01 | -0.01 | 0.00 | -0.00 | 0.00 |
|  | (0.01) | (0.01) | (0.01) | (0.01) | (0.01) | (0.00) | (0.01) |
| Age | -0.00 | -0.00 | -0.00 | 0.00 | -0.00 | -0.00 | -0.00 \*\* |
|  | (0.00) | (0.00) | (0.00) | (0.00) | (0.00) | (0.00) | (0.00) |
| DatasetCOBRE | -0.00 | 0.02 | -0.01 | 0.03 | 0.02 | -0.01 | 0.01 |
|  | (0.02) | (0.04) | (0.02) | (0.02) | (0.02) | (0.01) | (0.02) |
| DatasetHCP | -0.01 | -0.02 | -0.03 | 0.02 | -0.05 \*\* | 0.00 | -0.03 |
|  | (0.02) | (0.04) | (0.02) | (0.02) | (0.02) | (0.01) | (0.02) |
| CPZ\_eqiuv | -0.00 | 0.00 | -0.00 | -0.00 | 0.00 | 0.00 | 0.00 |
|  | (0.00) | (0.00) | (0.00) | (0.00) | (0.00) | (0.00) | (0.00) |
| CCBN:MCBN | -0.37 | -1.19 | -0.24 | -0.37 | -0.07 | 0.27 | -0.60 |
|  | (0.36) | (0.64) | (0.37) | (0.31) | (0.31) | (0.25) | (0.33) |
| CCBN:GroupCHR | 0.16 | -0.75 | -0.18 | -0.36 | -0.10 | 0.38 | -0.54 |
|  | (0.43) | (0.76) | (0.44) | (0.36) | (0.37) | (0.30) | (0.39) |
| CCBN:GroupECP | 0.15 | -0.27 | 1.12 \* | -0.01 | 1.06 \* | 1.10 \*\* | 0.15 |
|  | (0.52) | (0.93) | (0.54) | (0.45) | (0.46) | (0.36) | (0.48) |
| CCBN:GroupCP | 0.60 | -0.04 | -0.80 | -0.28 | 0.65 | 0.35 | 0.15 |
|  | (0.73) | (1.31) | (0.75) | (0.63) | (0.64) | (0.51) | (0.68) |
| MCBN:GroupCHR | -0.02 | -0.75 | -0.13 | -0.45 | -0.10 | 0.31 | -0.62 |
|  | (0.45) | (0.80) | (0.46) | (0.38) | (0.39) | (0.31) | (0.41) |
| MCBN:GroupECP | 0.48 | -0.05 | 1.12 | 0.21 | 1.26 \* | 1.29 \*\* | 0.36 |
|  | (0.58) | (1.03) | (0.59) | (0.49) | (0.51) | (0.40) | (0.53) |
| MCBN:GroupCP | 0.52 | -0.22 | -1.05 | -0.13 | 0.80 | 0.34 | 0.18 |
|  | (0.76) | (1.36) | (0.78) | (0.65) | (0.67) | (0.53) | (0.70) |
| CCBN:MCBN:GroupCHR | -0.04 | 1.29 | 0.18 | 0.65 | 0.17 | -0.58 | 0.93 |
|  | (0.64) | (1.14) | (0.66) | (0.55) | (0.56) | (0.45) | (0.59) |
| CCBN:MCBN:GroupECP | -0.63 | 0.45 | -1.50 | -0.27 | -1.81 \* | -1.76 \*\* | -0.43 |
|  | (0.80) | (1.44) | (0.83) | (0.69) | (0.71) | (0.56) | (0.74) |
| CCBN:MCBN:GroupCP | -0.96 | 0.21 | 1.45 | 0.07 | -0.98 | -0.53 | -0.19 |
|  | (1.07) | (1.91) | (1.10) | (0.91) | (0.94) | (0.74) | (0.98) |
| N | 438 | 438 | 438 | 438 | 437 | 438 | 438 |
| R2 | 0.08 | 0.08 | 0.08 | 0.11 | 0.14 | 0.08 | 0.17 |
| \*\*\* p < 0.001; \*\* p < 0.01; \* p < 0.05. | | | | | | | |

### Predicting Symptoms - Across All Groups (incl. HC)

#### PANSS-P

```
PANSS_P_all <- lm(PANSS.P ~ CCBN + MCBN + Alcohol + Marj + Age + Dataset + CPZ_eqiuv, data = dataset)
summary(PANSS_P_all)
```

```
## 
## Call:
## lm(formula = PANSS.P ~ CCBN + MCBN + Alcohol + Marj + Age + Dataset + 
##     CPZ_eqiuv, data = dataset)
## 
## Residuals:
##     Min      1Q  Median      3Q     Max 
## -19.849  -4.932  -1.623   4.542  21.686 
## 
## Coefficients:
##               Estimate Std. Error t value Pr(>|t|)    
## (Intercept) 11.3082808  4.4261874   2.555   0.0112 *  
## CCBN        -3.1546793  4.9056537  -0.643   0.5207    
## MCBN        -3.9012181  2.8399059  -1.374   0.1707    
## Alcohol      0.1178105  0.6466474   0.182   0.8556    
## Marj         0.7670158  0.5643716   1.359   0.1753    
## Age         -0.0525154  0.0399089  -1.316   0.1894    
## DatasetHCP   6.5877321  1.0742685   6.132 3.22e-09 ***
## CPZ_eqiuv    0.0037681  0.0004802   7.846 1.13e-13 ***
## ---
## Signif. codes:  0 '***' 0.001 '**' 0.01 '*' 0.05 '.' 0.1 ' ' 1
## 
## Residual standard error: 6.281 on 259 degrees of freedom
##   (181 observations deleted due to missingness)
## Multiple R-squared:  0.3689, Adjusted R-squared:  0.3518 
## F-statistic: 21.62 on 7 and 259 DF,  p-value: < 2.2e-16
```

```
PANSS_P_CCBN_all_plot <- effect_plot(PANSS_P_all, pred = CCBN, interval = TRUE, partial.residuals = TRUE)
PANSS_P_CCBN_all_plot + ylab("PANSS-P") + theme(text = element_text(size = 24), panel.border = element_rect(colour = "black", fill=NA, size=1))
```

```
PANSS_P_MCBN_all_plot <- effect_plot(PANSS_P_all, pred = MCBN, interval = TRUE, partial.residuals = TRUE)
PANSS_P_MCBN_all_plot + ylab("PANSS-P") + theme(text = element_text(size = 24), panel.border = element_rect(colour = "black", fill=NA, size=1))
```

#### PANSS-N

```
PANSS_N_all <- lm(PANSS.N ~ CCBN + MCBN + Alcohol + Marj + Age + Dataset + CPZ_eqiuv, data = dataset)
summary(PANSS_N_all)
```

```
## 
## Call:
## lm(formula = PANSS.N ~ CCBN + MCBN + Alcohol + Marj + Age + Dataset + 
##     CPZ_eqiuv, data = dataset)
## 
## Residuals:
##     Min      1Q  Median      3Q     Max 
## -23.455  -5.031  -2.076   4.332  17.997 
## 
## Coefficients:
##               Estimate Std. Error t value Pr(>|t|)    
## (Intercept) 16.5561307  4.7776163   3.465 0.000620 ***
## CCBN        -4.5110254  5.2951511  -0.852 0.395048    
## MCBN        -7.1161992  3.0653877  -2.321 0.021038 *  
## Alcohol     -0.6476066  0.6979896  -0.928 0.354366    
## Marj         0.6775389  0.6091814   1.112 0.267078    
## Age         -0.0734601  0.0430776  -1.705 0.089338 .  
## DatasetHCP   4.4557512  1.1595629   3.843 0.000153 ***
## CPZ_eqiuv    0.0036446  0.0005184   7.031 1.82e-11 ***
## ---
## Signif. codes:  0 '***' 0.001 '**' 0.01 '*' 0.05 '.' 0.1 ' ' 1
## 
## Residual standard error: 6.78 on 259 degrees of freedom
##   (181 observations deleted due to missingness)
## Multiple R-squared:  0.2922, Adjusted R-squared:  0.2731 
## F-statistic: 15.27 on 7 and 259 DF,  p-value: < 2.2e-16
```

```
PANSS_N_CCBN_all_plot <- effect_plot(PANSS_N_all, pred = CCBN, interval = TRUE, partial.residuals = TRUE)
PANSS_N_CCBN_all_plot + ylab("PANSS-N") + theme(text = element_text(size = 24), panel.border = element_rect(colour = "black", fill=NA, size=1))
```

```
PANSS_N_MCBN_all_plot <- effect_plot(PANSS_N_all, pred = MCBN, interval = TRUE, partial.residuals = TRUE)
PANSS_N_MCBN_all_plot + ylab("PANSS-N") + theme(text = element_text(size = 24), panel.border = element_rect(colour = "black", fill=NA, size=1))
```

#### PANSS-Cog

```
PANSS_C_all <- lm(PANSS.Cog ~ CCBN + MCBN + Alcohol + Marj + Age + CPZ_eqiuv, data = dataset)
summary(PANSS_C_all)
```

```
## 
## Call:
## lm(formula = PANSS.Cog ~ CCBN + MCBN + Alcohol + Marj + Age + 
##     CPZ_eqiuv, data = dataset)
## 
## Residuals:
##     Min      1Q  Median      3Q     Max 
## -9.6882 -1.2537 -0.2705  1.3625  6.0455 
## 
## Coefficients:
##              Estimate Std. Error t value Pr(>|t|)    
## (Intercept)  5.027497   2.845298   1.767  0.08009 .  
## CCBN         9.169216   3.131570   2.928  0.00417 ** 
## MCBN        -2.307249   1.967310  -1.173  0.24348    
## Alcohol     -0.089567   0.399696  -0.224  0.82312    
## Marj         0.019725   0.296514   0.067  0.94709    
## Age          0.008272   0.072483   0.114  0.90935    
## CPZ_eqiuv    0.004868   0.001187   4.102 8.01e-05 ***
## ---
## Signif. codes:  0 '***' 0.001 '**' 0.01 '*' 0.05 '.' 0.1 ' ' 1
## 
## Residual standard error: 2.711 on 107 degrees of freedom
##   (334 observations deleted due to missingness)
## Multiple R-squared:  0.2373, Adjusted R-squared:  0.1945 
## F-statistic: 5.548 on 6 and 107 DF,  p-value: 4.871e-05
```

```
PANSS_Cog_CCBN_all_plot <- effect_plot(PANSS_C_all, pred = CCBN, interval = TRUE, partial.residuals = TRUE)
PANSS_Cog_CCBN_all_plot + ylab("PANSS-Cog") + theme(text = element_text(size = 24), panel.border = element_rect(colour = "black", fill=NA, size=1))
```

```
PANSS_Cog_MCBN_all_plot <- effect_plot(PANSS_C_all, pred = MCBN, interval = TRUE, partial.residuals = TRUE)
PANSS_Cog_MCBN_all_plot + ylab("PANSS-Cog") + theme(text = element_text(size = 24), panel.border = element_rect(colour = "black", fill=NA, size=1))
```

#### SIPS-P

```
SIPS_P_all <- lm(SIPS.P ~ CCBN + MCBN + Alcohol + Marj + Age, data = dataset)
summary(SIPS_P_all)
```

```
## 
## Call:
## lm(formula = SIPS.P ~ CCBN + MCBN + Alcohol + Marj + Age, data = dataset)
## 
## Residuals:
##    Min     1Q Median     3Q    Max 
## -9.427 -4.712 -1.031  3.954 17.805 
## 
## Coefficients:
##             Estimate Std. Error t value Pr(>|t|)    
## (Intercept)  -1.5842     5.5714  -0.284   0.7765    
## CCBN         -6.3237     4.5532  -1.389   0.1668    
## MCBN          3.1305     3.0702   1.020   0.3094    
## Alcohol       4.2321     1.0379   4.078 7.06e-05 ***
## Marj          7.4725     1.7723   4.216 4.08e-05 ***
## Age           0.4717     0.1948   2.422   0.0165 *  
## ---
## Signif. codes:  0 '***' 0.001 '**' 0.01 '*' 0.05 '.' 0.1 ' ' 1
## 
## Residual standard error: 5.817 on 165 degrees of freedom
##   (277 observations deleted due to missingness)
## Multiple R-squared:  0.2487, Adjusted R-squared:  0.226 
## F-statistic: 10.93 on 5 and 165 DF,  p-value: 4.273e-09
```

```
SIPS_P_CCBN_all_plot <- effect_plot(SIPS_P_all, pred = CCBN, interval = TRUE, partial.residuals = TRUE)
SIPS_P_CCBN_all_plot + ylab("SIPS-P") + theme(text = element_text(size = 24), panel.border = element_rect(colour = "black", fill=NA, size=1))
```

```
SIPS_P_MCBN_all_plot <- effect_plot(SIPS_P_all, pred = MCBN, interval = TRUE, partial.residuals = TRUE)
SIPS_P_MCBN_all_plot + ylab("SIPS-P") + theme(text = element_text(size = 24), panel.border = element_rect(colour = "black", fill=NA, size=1))
```

#### SIPS-N

```
SIPS_N_all <- lm(SIPS.N ~ CCBN + MCBN + Alcohol + Marj + Age, data = dataset)
summary(SIPS_N_all)
```

```
## 
## Call:
## lm(formula = SIPS.N ~ CCBN + MCBN + Alcohol + Marj + Age, data = dataset)
## 
## Residuals:
##     Min      1Q  Median      3Q     Max 
## -12.322  -3.336  -1.926   2.181  19.398 
## 
## Coefficients:
##             Estimate Std. Error t value Pr(>|t|)    
## (Intercept)  -0.2772     5.2963  -0.052 0.958316    
## CCBN         -5.2717     4.3284  -1.218 0.224982    
## MCBN          4.8363     2.9186   1.657 0.099411 .  
## Alcohol       3.7114     0.9866   3.762 0.000234 ***
## Marj          7.3051     1.6848   4.336 2.52e-05 ***
## Age           0.2067     0.1852   1.116 0.265949    
## ---
## Signif. codes:  0 '***' 0.001 '**' 0.01 '*' 0.05 '.' 0.1 ' ' 1
## 
## Residual standard error: 5.53 on 165 degrees of freedom
##   (277 observations deleted due to missingness)
## Multiple R-squared:  0.2113, Adjusted R-squared:  0.1874 
## F-statistic: 8.843 on 5 and 165 DF,  p-value: 1.866e-07
```

```
SIPS_N_CCBN_all_plot <- effect_plot(SIPS_N_all, pred = CCBN, interval = TRUE, partial.residuals = TRUE)
SIPS_N_CCBN_all_plot + ylab("SIPS-N") + theme(text = element_text(size = 24), panel.border = element_rect(colour = "black", fill=NA, size=1))
```

```
SIPS_N_MCBN_all_plot <- effect_plot(SIPS_N_all, pred = MCBN, interval = TRUE, partial.residuals = TRUE)
SIPS_N_MCBN_all_plot + ylab("SIPS-N") + theme(text = element_text(size = 24), panel.border = element_rect(colour = "black", fill=NA, size=1))
```

#### Combined table

```
export_summs(PANSS_P_all, PANSS_N_all, PANSS_C_all, SIPS_P_all, SIPS_N_all, to.file = "docx", file.name = "models_symptoms.docx", model.names = c("PANSS-P", "PANSS-N", "PANSS-C", "SIPS-P", "SIPS-N"))
```

|  | PANSS-P | PANSS-N | PANSS-C | SIPS-P | SIPS-N |
| --- | --- | --- | --- | --- | --- |
| (Intercept) | 11.31 \* | 16.56 \*\*\* | 5.03 | -1.58 | -0.28 |
|  | (4.43) | (4.78) | (2.85) | (5.57) | (5.30) |
| CCBN | -3.15 | -4.51 | 9.17 \*\* | -6.32 | -5.27 |
|  | (4.91) | (5.30) | (3.13) | (4.55) | (4.33) |
| MCBN | -3.90 | -7.12 \* | -2.31 | 3.13 | 4.84 |
|  | (2.84) | (3.07) | (1.97) | (3.07) | (2.92) |
| Alcohol | 0.12 | -0.65 | -0.09 | 4.23 \*\*\* | 3.71 \*\*\* |
|  | (0.65) | (0.70) | (0.40) | (1.04) | (0.99) |
| Marj | 0.77 | 0.68 | 0.02 | 7.47 \*\*\* | 7.31 \*\*\* |
|  | (0.56) | (0.61) | (0.30) | (1.77) | (1.68) |
| Age | -0.05 | -0.07 | 0.01 | 0.47 \* | 0.21 |
|  | (0.04) | (0.04) | (0.07) | (0.19) | (0.19) |
| DatasetHCP | 6.59 \*\*\* | 4.46 \*\*\* |  |  |  |
|  | (1.07) | (1.16) |  |  |  |
| CPZ\_eqiuv | 0.00 \*\*\* | 0.00 \*\*\* | 0.00 \*\*\* |  |  |
|  | (0.00) | (0.00) | (0.00) |  |  |
| N | 267 | 267 | 114 | 171 | 171 |
| R2 | 0.37 | 0.29 | 0.24 | 0.25 | 0.21 |
| \*\*\* p < 0.001; \*\* p < 0.01; \* p < 0.05. | | | | | |
